# Metrics for studying the porous void space of packed particles

**DOI:** 10.1101/2025.07.23.666315

**Authors:** Lindsay Riley, Emma Lee, Peter Cheng, Daniel Adrianzen Alvarez, Tatiana Segura

**Affiliations:** Department of Biomedical Engineering, Duke University, Durham, North Carolina, 27708, USA; Yale University, New Haven, Connecticut, 06520, USA; Ninjabyte Computing, Sunnyvale, California, 94086, USA; Department of Medicine, Neurology, Dermatology, Duke University, Durham, North Carolina, 27708, USA

**Author notes:** equal contribution (Lindsay Riley, Emma Lee).

## Abstract

Characterizing porosity in packed particle assemblies is a complex task that requires advanced analytical tools. We present a visually rich and extensive library of global, pore-based, and other metrics for analyzing features of porosity in such assemblies. Our library includes over 25 descriptors of “3D pores” that are identified using our LOVAMAP software. By applying our metrics to a set of simulated packings that vary by particle size, shape, and stiffness, we reveal predictable relationships between particle and void space characteristics. We identify two fundamental parameters of a monodisperse particle system – particle diameter (*δ*) and void volume fraction (*ϕ*) – that govern several void space features, such as the total number of bottlenecks (i.e., doors between pores), the median value of the largest enclosed sphere across all pores in a packing, and the fraction of reaction-center “hotspots.” Through regression analyses on transformations of *δ* and *ϕ*, we quantify multiple packing-descriptor relationships, demonstrating, for example, that packing properties scale linearly with the median values of length-based descriptors across assemblies. We further introduce approaches for computing the number of vertices, edges, and faces of 3D pores, allowing for approximation to simpler polyhedra. Additional metrics explore surface entrances into the particle scaffold, traversable paths through the void space, and size-based accessibility. Together, these descriptors, which have been bundled into LOVAMAP, offer new insights into particle-pore architecture and spatial organization.

## Introduction

Porous materials are ubiquitous in natural phenomena and engineered constructs across disciplines, and for many porous systems, the structure of the void space plays a critical role. Such void space is a network of interconnected pockets or channels that enable processes like mass transport, fluid flow, and object migration, where changes in local architecture can have notable impact on material behavior and performance(2-4). In subsurface hydrology and petroleum engineering, understanding pore space distributions has been instrumental in predicting reservoir performance(5), physical properties of rocks(6), digital modeling(7), and optimizing fluid extraction strategies. Studies of natural sedimentary formations have shown that increased tortuosity in pore networks leads to lower permeability, directly impacting hydrocarbon recovery rates(8). Similarly, in bioengineering, granular porous materials serve as platforms for cell culture(9), bioprinting(10, 11), drug delivery(12), and tissue engineering(13-16), where the arrangement and connectivity of porosity directly impact biological integration and nutrient transport. In these biomaterial scaffolds, experimental work has shown that optimized pore architectures enhance tissue regeneration outcomes(17). These examples highlight the importance of studying porosity and developing metrics to characterize it.

Characterizing granular void space architecture is challenging due to its continuous, tortuous morphology, whose irregular surfaces make three-dimensional (3D) analysis particularly complex. A traditional simplification involves examining two-dimensional (2D) slice data, which can provide useful information despite inherent limitations(18, 19). However, advancements in imaging and analytical tools are enabling the transition from 2D approximations to more accurate analyses of 3D reconstructions(20, 21).

Regardless of dimensionality, reporting the proportion of void space volume relative to total volume of a material offers a straightforward way to ignore the geometric complexity of the void space. For decades, this single-scalar metric has served to characterize overall porosity and has facilitated easy-to-interpret comparisons across constructs(2, 22). A key limitation of global descriptors like void volume fraction, however, is their inability to capture detailed information about local architecture or the heterogeneity of the porous network.

To this end, methodologies have been developed to segment the void space into subspaces(23-26). Ideally, each subspace delineates the widest local region – termed a pore. Pores are often either physically bounded by the solid phase or conceptually bounded by narrow portions of the empty phase, referred to as doors, necks, throats, or bottlenecks. Partitioning the continuous void space into pores separated by doors not only enables analysis of individual pores but also facilitates the study of connectivity between them. Connectivity influences how fluids, solutes, cells, and other entities percolate through the network, making it a frequently-examined property in porous materials research(27, 28).

Porosity itself can be static or dynamic depending on the structure of the solid phase. For example, the pore architecture is fixed in samples of sandstone and trabecular bone because the solid phase is immobile. In contrast, the solid phase of packed granular material is mobile because particles can shift, which allows for dynamic porosity between touching particles. Pore geometry in granular material is therefore directly influenced by particle properties like packing fraction, configuration, size, shape, stiffness, roughness, and polydispersity. Previous studies have examined the relationship between particle composition and void volume fraction, uncovering several fundamental insights. For instance, closely-packed monodisperse rigid spheres yield a void volume fraction of approximately 0.36, a value that is scale-independent across different particle diameters(29, 30). As another example, mixing rigid spheres of two different diameters yields the tightest packing when the ratio of small-to-large particles (by volume) is approximately 35:65(22, 31). Weeks et al. 2024 expand upon this field of study by proposing a linear regression equation to estimate the void volume fraction of polydisperse spherical packings as a function of polydispersity (dependent on radius mean and standard deviation) and skewness, for large enough skew(32).

In Riley et al. 2023, we pivot from void volume fraction to explore relationships between particles and pores. This was accomplished by developing a software (termed LOVAMAP) that segments void space into 3D pores within packed particles(1). For the purpose of orienting the reader to relevant terminology, we briefly review the approach before highlighting results. LOVAMAP computes the Euclidean distance transform (EDT) of the void space for each particle in order to identify subtypes of the medial axis – namely 2D ridges, one-dimensional (1D) ridges, and peaks, where 2D ridges are surfaces equidistant to two particles, 1D ridges lie at intersections of 2D ridges and are curves equidistant to 3 or more particles, and peaks lie at intersections of 1D ridges and are local center points in space. Peaks and 1D ridges form a graph of nodes and edges, respectively, which we refer to as the backbone of the void space. LOVAMAP segments this graph to form the backbone of each pore according to two principles: peaks that belong to the same open space remain connected, and pore delineation occurs at the narrowest point along 1D ridges. The backbones are then used to construct the complete 3D pores (or simply, pores). For each pore, LOVAMAP computes basic metrics including volume, length, and largest enclosed sphere. In our previous publication, we report empirical equations through curve fitting that describe the relationship between particle properties and the total number and median size of pores across a range of simulated particle assemblies in random packing. We also confirm that pores enclosed by four particles are the most prevalent pore type within each packing. For packings comprising monodisperse rigid spheres, we reveal that the volume of the largest possible pores enclosed by four particles rarely overlaps with the volume of the smallest pores enclosed by five particles; the trend continues for pores surrounded by five, six, etc. particles. Our findings uncover patterns in the void space that are otherwise hidden when restricted to particle-centric analysis.

In the current work, we report new descriptors that expand upon pore analysis of packed particles and extend beyond it. Our additional pore descriptors provide information about pore size, shape, directionality, and particle-surface influence on the pore space. We also include descriptors that convey information about connectivity, both within the void space and among the particles. Since objects, such as infiltrating cells, can enter or exit packed particles at the surface of the scaffold, we offer descriptors that address these openings on the surface. We introduce two additional concepts: paths and available regions. Paths represent the shortest distance from the center of the void space to the exit points of the structure, while available regions indicate which areas of the void space can accommodate objects of a specified size. Lastly, the collective concepts covered by LOVAMAP contribute to a set of global descriptors that provide single-value characterizations of a packing. A complete list of the descriptors covered in this article are provided in Supplementary Tables 1, 2, and 3. As in the development of LOVAMAP, we use simulations of monodisperse packed particles to study our descriptors and reveal relationships between particle properties and porosity (Figure 1a).

**Fig. 1.**
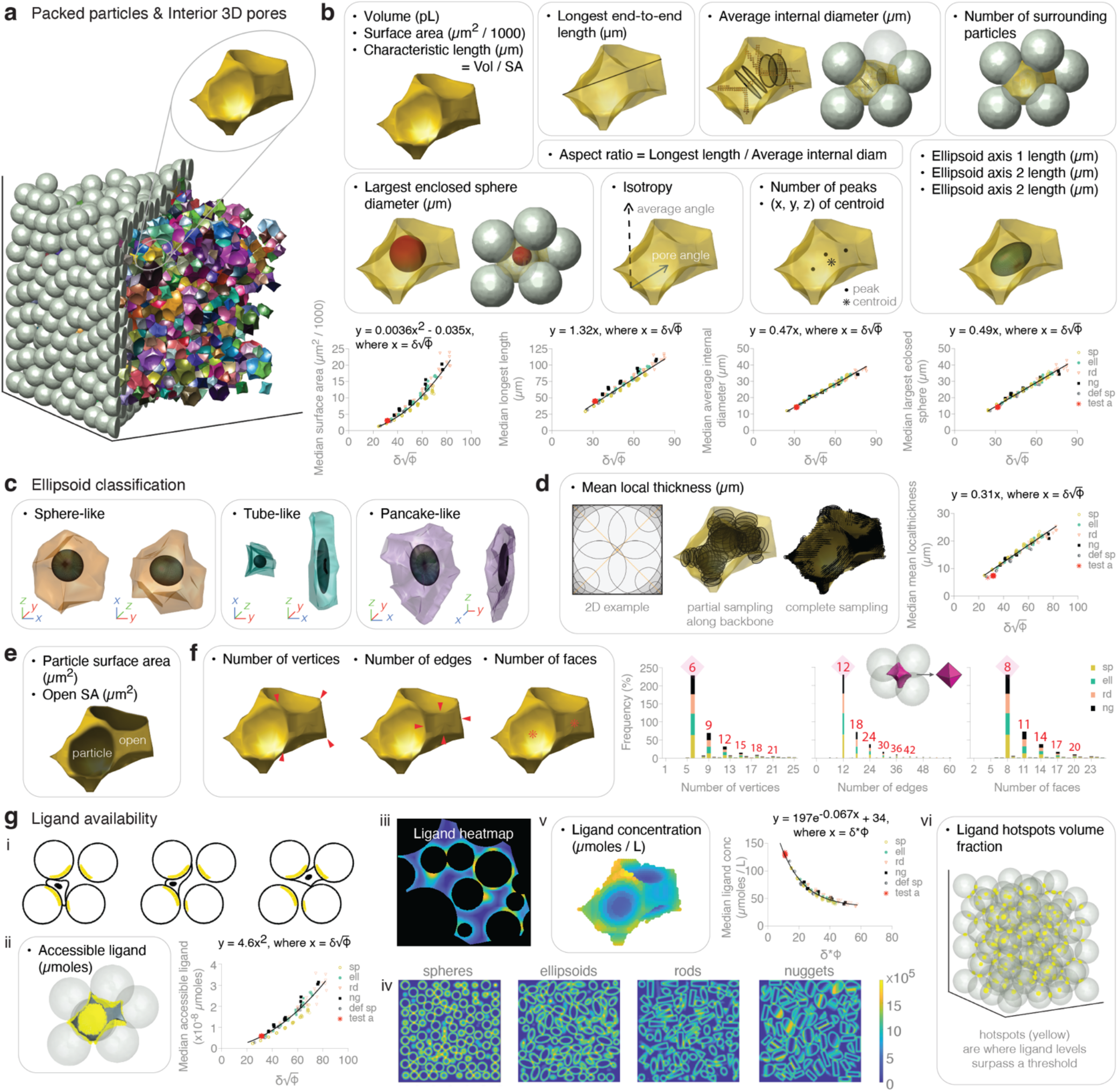
3D-pore descriptors. **a**, Sample packed particle assembly (left, gray) with corresponding interior 3D pores (right, distinct colors), highlighting a yellow pore. **b**, 3D-pore descriptors introduced in Riley, et al. 2023(1). For Average Internal Diameter, sample diameters (black circles) are displayed along central pore backbone. **c**, Using pore ellipsoids derived from principal component analysis to identify three pore-shape categories. **d**, Mean local thickness (µm) of pores; 2D example demonstrates calculation for a square, where thickness measurements along backbone (orange) are represented by black circles. Equation predicts the median mean local thickness for monodisperse packings given δ and ϕ. **e**, Particle surface area (µm^2^) and open surface area (µm^2^) of pores. **f**, Number of vertices, edges, and faces of pores with corresponding distribution plots of monodisperse 60 µm diameter particles. For pore images, annotating examples of each element (red markers); for plots, labeling frequency spikes (red numbers). **g**, Studying concepts of protein (ligand) attached to the surface of particles. (i) Schematic of a migrating cell interacting with ligand molecules on the surface of particles that surround it. (ii) Accessible ligand (µmoles) of pores, where yellow indicates ligand on the surface of particles surrounding the pore. Note the transparent particle projecting toward the reader at the center of the cluster. Equation predicts the median accessible ligand for monodisperse packings given δ and ϕ. (iii) 2D slice of a ligand heatmap representing the concentration of ligand as sensed by a cell migrating through the void space. Non-void regions are masked in black to highlight ligand activity within the void space. Color key shown in the subsequent subpanel. (iv) 2D slices of ligand heatmaps for packed spheres, ellipsoids, rods, and nuggets. (v) Ligand concentration (µmoles / L) of pores. Equation predicts the median ligand concentration for monodisperse packings given δ and ϕ. (vi) Ligand hotspots volume fraction of scaffolds; yellow regions indicate highest ligand levels. δ, particle diameter; ϕ, void volume fraction; sp, spheres; ell, ellipsoids; rd, rods; ng, nuggets; def sp, deformable spheres; test a, unbiased estimator of softer deformable spheres with δ = 100, λ= 400, μ = 50. N = 10 domains (sp, def sp, test a) and N = 4 – 5 domains (ell, rd, ng).

## Results

### 3D pore size, shape, and directionality

Pore descriptors provide detailed information about the pockets of open space in a packed particle assembly. We first focus on interior pores that do not extend to the surface of the void space. Figure 1b provides visual representations of descriptors that were introduced in Riley et al. 2023(1). Surface area and characteristic length (volume per surface area) of pores have been included. Most of the descriptors are self-explanatory, but briefly, average internal diameter refers to the average spatial width along the backbone of the pore. This value is used as the width measurement in our computation of 3D aspect ratio, where height is taken as the longest end-to-end length of the pore. Our isotropy measurement indicates how aligned a pore is to the average pore orientation, ranging from orthogonal (value of 0) to parallel (value of 1). If all pores have isotropy values close to 1, this suggests a high degree of anisotropy in the system.

As a resource for approximating select descriptors without running LOVAMAP, we fit empirical equations to descriptor data of simulated particle domains. Domains comprise randomly-packed monodisperse spheres, ellipsoids (prolate spheroids), rods, nuggets, and deformable spheres (Lamé parameters λ = 1,000, μ = 250)(1). Each equation incorporates a transformation of the average particle diameter, *δ*, (in micrometers) and void volume fraction, *ϕ*, of the packing – two parameters that govern the structure of the void space. Below are our results for predicting the median pore value of surface area (µm^2^ / 1000), longest length (µm), average internal diameter (µm), and largest enclosed sphere (µm), respectively, for a given packing with known *δ* and *ϕ* (Figure 1b):

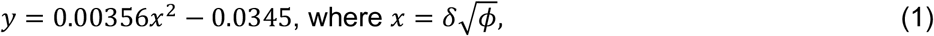

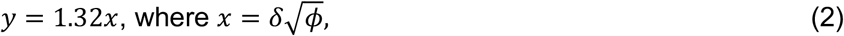

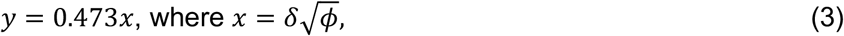

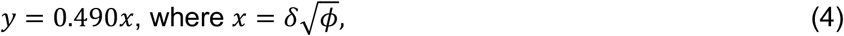

(R^2^ = 0.9381, 0.9492, 0.9862, and 0.9802, respectively). For non-spherical particles, *δ* is the diameter of a sphere with equivalent volume. In our plots, we include a test dataset (test a) comprising softer deformable spheres (*δ* = 100, Lamé parameters λ = 400, μ = 50) to serve as an unbiased estimator. The normalized root-mean-squared error (NRMSE) equals 0.0355, 0.00450, 0.00331, 0.00805, respectively, which suggests good model fits. Our results reveal a tight relationship between packing properties and the median descriptor value of pores in the packing.

We next introduce new pore descriptors that provide both spatial metrics and biological relevancy. The first descriptor is categorical, using principal-component-analysis (PCA) ellipsoids (Figure 1b) to classify pores as 1) sphere-like, 2) tube-like, or 3) pancake-like (Figure 1c). The latter two shapes are more frequently observed in packings of deformable or non-spherical particles or in those confined within a spherical container.

Mean local thickness is a measurement adapted from Hildebrand and Ruegsegger(33) that uses enclosed spheres to assign local thickness measurements throughout the pore, which are then averaged (Figure 1d). For a given point in space, its local thickness equals the diameter of the largest enclosed sphere that contains the point. As a simple 2D reference, a circle of diameter *δ* has a mean local thickness equal to *δ*, whereas a square of length *δ* has a smaller mean local thickness because the local thickness values decrease toward the corners, reducing the average (Figure 1d, Supplementary Note 1). With curve-fitting, we estimate the median mean local thickness of pores (R^2^ = 0.9677):

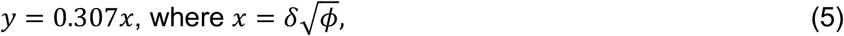

where *δ* and *ϕ* are defined above. A NRMSE of 0.0715 indicates good model agreement. As with Equations 2 – 4, we observe a descriptor with units of length that follows a linear relationship when plotted against the transformation 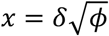 (Figure 1d).

In Riley et al. 2023, we study pores according to the number of surrounding particles (Figure 1b). Here, we take this further by quantifying total surface area contributed by surrounding particles, as well as total surface area contributed by open (non-particle) transitions into neighboring pores (Figure 1e), providing insight into how tightly packed particles are around a pore (Supplementary Figure 1).

LOVAMAP utilizes 1D-and 2D-ridge information to identify the vertices, edges, and faces of pores (Figure 1f, Supplementary Note 2). Defining such features are useful for approximating pores to genus-0 polyhedra, i.e., polyhedra that lack holes. In addition, edge-to-vertex ratio can be viewed as a complexity metric for a given pore, where more edges per vertex indicates a more complex shape.

When we plot histograms of vertex, edge, and face data for pores in scaffolds comprising 60 µm diameter rigid spheres, ellipsoids, rods, or nuggets, a combinatorial pattern of spikes emerges (red annotations in Figure 1f) – one that can be explained by the Euler characteristic(34). For polyhedra of genus-0, the Euler characteristic formula simplifies to *V – E + F =* 2, where *V, E*, and *F* are the number of vertices, edges, and faces, respectively. This relationship enforces a tight coupling of *V, E*, and *F*, which is reflected in the distinct binning patterns of the histograms. As a concrete example, consider the smallest and most common pore type formed by four packed particles enclosing a single space(1). This pore is represented by the tallest bin in each of our histogram plots: *V =* 6, *E =* 12, and *F =* 8. Indeed, the Euler characteristic of this pore type is 2, and its *V, E, F* counts match that of an octahedron. A side-by-side comparison of pore and octahedron is shown as an inset in Figure 1f. Our results from Figure 1f suggest that the geometry of simpler 3D pores is governed by the same topological relationship as genus-0 polyhedra.

### Particle surface ligand availability

Several applications using packed particles rely on interactions at the interface of particles and pores. We study this concept using a biological example of cells migrating between particles in a granular scaffold, where particles coated with integrin-binding ligands (e.g., RGD peptides) mediate cell adhesion (Figure 1g *i*)(35). To mimic this scenario, we assign a ligand count to voxels at the surface of each particle. This allows us to report the µmoles of ligand that are accessible within a given pore (Figure 2g *ii*). To predict median accessible ligand, we fit an exponential curve (R^2^ = 9113, NRMSE of 0.04):

**Fig. 2.**
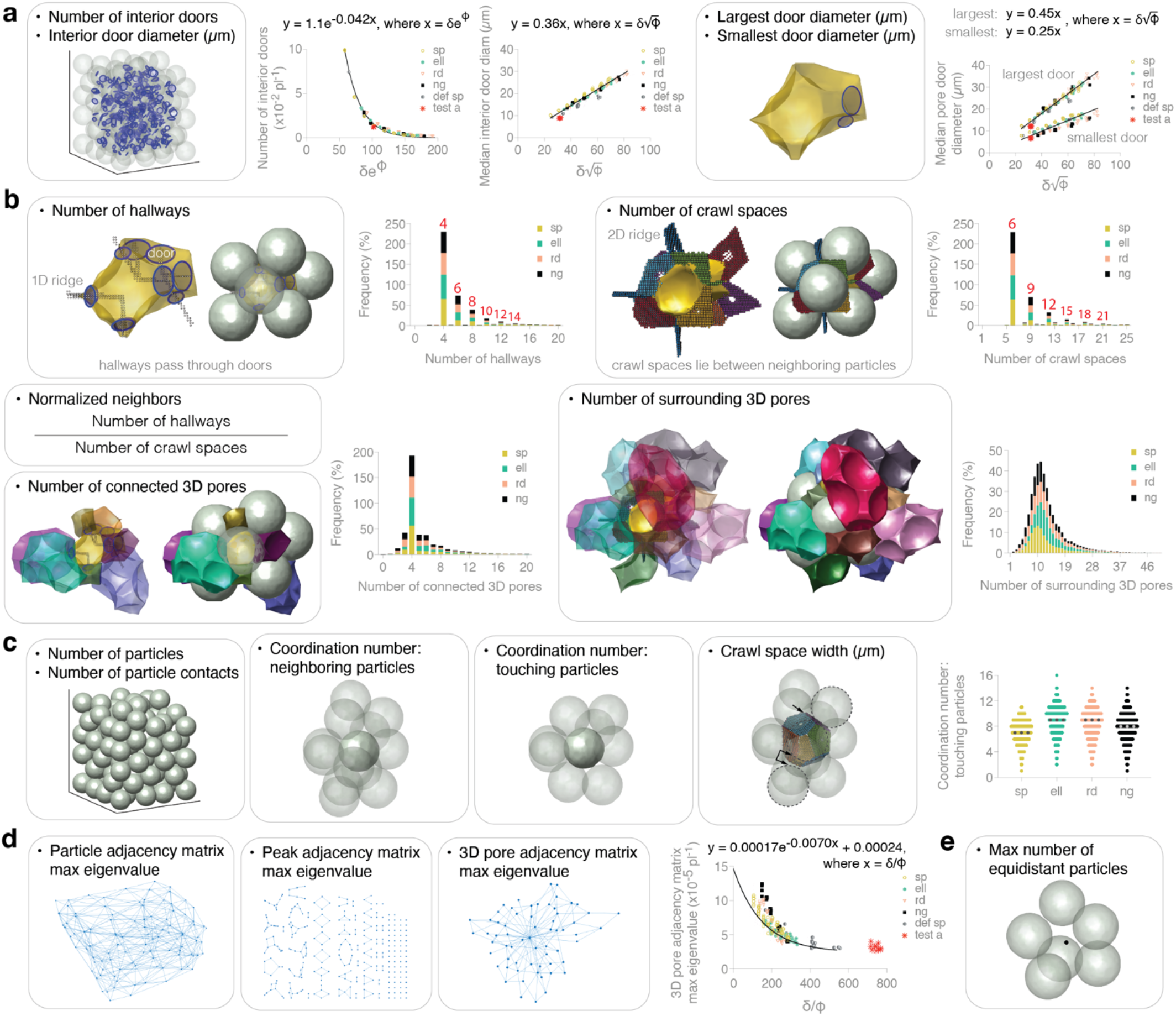
Connectivity descriptors of doors, pores, and particles. **a**, (left) Number and diameter (µm) of interior doors within a scaffold (doors represented as blue circles). Equations predict the total number of interior doors and the median internal door diameter for monodisperse packings given δ and ϕ. (right) Largest and smallest door diameter (µm) of pores. Equations predict the median of these values for monodisperse packings given δ and ϕ. **b**, Number of hallways, crawl spaces, connected 3D pores, and surrounding 3D pores of pores with corresponding distribution plots of monodisperse 60 µm diameter particles. Red numbers label frequency spikes. Definition of normalized neighbors of pores also displayed. Note: hallways exist along 1D ridges; crawl spaces exist along 2D ridges. **c**, Number of particles and particle contacts, coordination number (defined by neighboring particles vs. touching particles), and crawl space width between neighboring particles within a scaffold. Showing scatter plots of coordination number (by touching particles) for monodisperse 60 µm diameter particles. **d**, Maximum eigenvalue of adjacency matrices for scaffolds; showing particle, peak, and pore graph diagrams. Equation predicts the pore adjacency matrix eigenvalue for monodisperse packings given δ and ϕ. **e**, Maximum number of equidistant particles within a scaffold (centered around a peak), which is influenced by particle configuration and voxel resolution. δ, particle diameter; ϕ, void volume fraction; sp, spheres; ell, ellipsoids; rd, rods; ng, nuggets; def sp, deformable spheres; test a, unbiased estimator of softer deformable spheres with δ = 100, λ= 400, μ = 50. N = 10 domains (sp, def sp, test a) and N = 4 – 5 domains (ell, rd, ng).

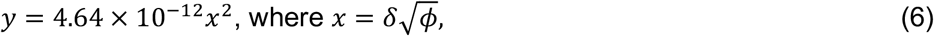

where *δ* and *ϕ* are as above (Figure 2g *ii*). Since accessible ligand is reported as an absolute quantity, we expect the value to increase as particle size increases.

We next transform the particle domain into a ligand heatmap that mimics diffusion of ligand into the void space. To the domain, we apply a spherical averaging-filter with diameter 10 µm (Figure 1g *iii*). The filter size is the approximate size of a cell, allowing the resulting heatmap to represent the ligand landscape as sensed by a migrating cell. Figure 1g *iv* shows 2D-slice samples of ligand heatmaps for various particle packings, where particle ligand is confined to a 4 µm-thick outer shell that represents the depth accessible by cells. With these heatmaps, we compute total ligand concentration of pores (in µmoles / L) (Figure 1g *v*). Pores formed by tightly packed particles will exhibit higher heatmap-derived ligand concentration relative to looser packing. To this end, we again incorporate void volume fraction into the transformation of our independent variable for predicting median pore ligand concentration of a scaffold (R^2^ = 0.9730):

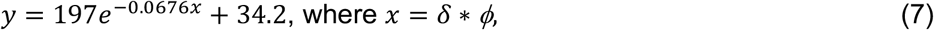

where *δ* is particle diameter and *ϕ* is void volume fraction as explained previously (Figure 1g *v*). Our test group with NRMSE of 0.000973 supports a robust model. Our heatmaps also allow us to identify ligand hotspots of particularly high ligand counts, which we report as a percentage of the total void space. The adjustable threshold for high ligand count was chosen at 90% of the maximum across all heatmaps in our study. A visual representation of ligand hotspots showcases the spatial distribution within a granular scaffold of single-species rigid monodisperse spheres, which correspond to particle-particle contact points (Figure 1g *vi*).

### Connectivity among pores and particles

Concepts of connectivity are integral to the structure and function of packed particle systems. Our discussion of connectivity begins with interior doors (displayed as blue circles), which are separation planes between pores and represent the narrowest circular cross section along the backbone of the void space. In other words, interior doors are bottlenecks of the porous space where flow and migration are most hindered. For this reason, we provide equations for the total number of interior doors per picoliter (R^2^ = 0.9888, NRMSE = 0.0462) and median door diameter (R^2^ = 0.9515, NRMSE = 0.0870) within a particle packing using the following transformations, respectively:

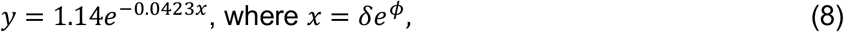

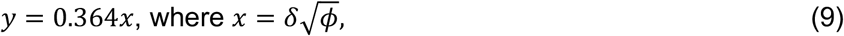

where *δ* and *ϕ* are as above (Figure 2a).

We also study doors on a per-pore basis, identifying the largest and the smallest door of a pore. The following equations can be used to approximate the median value of the largest (R^2^ = 0.9650, NRMSE = 0.0289) and smallest (R^2^ = 0.7177, NRMSE = 0.0253) door diameter (per pore) in a packing, respectively:

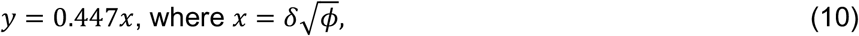

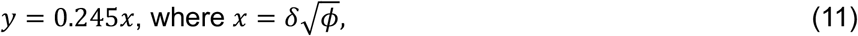

where *δ* is particle diameter and *ϕ* is void volume fraction (Figure 2a).

We define two types of connectivity between pores: 1) hallways, which pass through doors along 1D ridges, and 2) crawl spaces, which are between touching particles along 2D ridges (Figure 2b, Supplementary Note 2). LOVAMAP outputs the number of hallways exiting a pore normalized by the number of crawl spaces – a metric termed normalized neighbors – which alludes to particle compactness.

As with vertices, edges, and faces of pores, we recognize combinatorial spikes in the histogram plots of hallways and crawl spaces for pores (red annotations in Figure 2b, top). Our crawl-space results can be understood by viewing each 3D pore as empty space within a mini particle cluster. The tallest bin in the crawl-space plot refers to 4-particle clusters (i.e., pores surrounded by four particles), which is the most common pore type(1). The second tallest bin refers to 5-particle clusters, etc. In jammed systems of rigid particles, clusters are most commonly isostatic, i.e., the number of particle-particle contacts, *N*_c_, within a given cluster equals 3*n* − 6, where *n* = the number of particles(36). Since crawl spaces exist between particle-particle contacts, *N*_c_, approximates the number of crawl spaces of a pore that is surrounded by *n* particles.

For our hallway results, the binning pattern can be explained by recognizing that most hallways exiting smaller pores pass through three touching particles – referred to as triangular cliques – which form the simplest hallway type. Again viewing 3D pores as mini particle clusters, we empirically identify the number of triangular cliques, *N*_*T*_, for a particle cluster as 2*n* − 4, where *n* = the number of particles. Therefore, *N*_*T*_ approximates the number of hallways of a pore that is surrounded by *n* particles.

Pores connected to other pores through hallways are not always one-to-one, so for a given pore, we report the number of connected 3D pores that are accessible through hallways, i.e.,1D ridges (Figure 2b, bottom). Similarly, we report the number of surrounding 3D pores that are accessible through crawl spaces, i.e., 2D ridges (Figure 2b, bottom). Since 1D ridges lie at the intersection of 2D ridges, connected 3D pores are a subset of surrounding 3D pores. Not surprisingly, resulting histogram plots form smooth spreads relative to the hallways and crawl spaces plots that are constrained by scaling laws. These results emphasize the complexity of pore connectivity since multiple pores may be connected through the same crawl space and multiple hallways may span the same two pores.

Shifting our discussion to particles, we report classic metrics such as the total number of particles, the total number of particle-particle contacts (Supplementary Figure 2), and coordination number per particle (Figure 2c). Coordination number for a particle is reported in two ways: 1) neighboring particles, which refers to all particles that share a 2D ridge with the particle, and 2) touching particles, which only considers particles that are in physical contact with the particle (Figure 2c). Touching particles are a subset of neighboring particles.

As previously mentioned, crawl spaces refer to the space between two particles, and we report the widths of all crawl spaces in the packing, which equal 0 µm when particles are touching (Figure 2c). We show swarm plots of particle coordination numbers – specifically, touching particles – in scaffolds of 60 µm diameter rigid spheres, ellipsoids, rods, and nuggets (Figure 2c). Randomly packed frictionless spheres typically yield a median coordination number of six(37), but incorporation of friction in our simulations produces a slightly higher median of seven. As expected, particle coordination numbers for spheres do not exceed 12(37).

To capture other aspects of connectivity, we analyze adjacency matrices for three systems: particles, peaks, and pores (Figure 2d). Particle adjacency exists if two particles share a 2D ridge. Peak adjacency exists if the distance between two peaks is less than the sum of their EDT values, i.e., the peaks are part of the same open space. Pore adjacency exists if two pores are connected by a hallway. Using these objects as our nodes, we construct our adjacency matrices and report the maximum eigenvalues of the matrices, which is bounded below by the average number of edges per node and above by the maximum number of edges for any node. Focusing on 3D pore adjacency matrices, we plot the maximum eigenvalue for our scaffolds and fit the following curve:

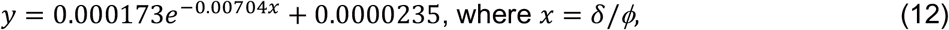

where *δ* and *ϕ* are particle diameter and void volume fraction, respectively (R^2^ = 0.7487, NRMSE = 0.0678). Lastly, for each particle packing, we output the maximum number of particles that are equidistant to a single peak – a metric that is influenced by both randomness and grid resolution (Figure 2e).

### Entering and exiting a packed particle scaffold

We next address pores that extend to the outside of the packing, termed surface 3D pores. As with interior pores, surface 3D pores are depicted as solid, colorful objects that represent empty space (Figure 3a). We define an imaginary boundary at the surface of the void space through which objects pass to enter or exit the scaffold between particles. Mathematically, this boundary is the non-particle surface of the convex hull of particle centers. To form surface 3D pores, LOVAMAP computes the EDT along the surface (Figure 3b, left) as a first step toward identifying the natural openings into and out of the scaffold (termed surface 2D pores) (Figure 3b, right). Interior pores that share the same surface 2D pore are then combined to form the final surface 3D pore(1). This approach can produce very large surface 3D pores when multiple large openings connect pores within (Supplementary Figure 3). To quantify this point, LOVAMAP outputs the maximum number of peaks contained within a single surface 3D pore, which captures surface pore size and complexity. Figure 3c showcases a particularly large and interconnected surface pore that can be attributed to large particle size relative to container size, which contrasts the surface pores shown in Figure 3a.

**Fig. 3.**
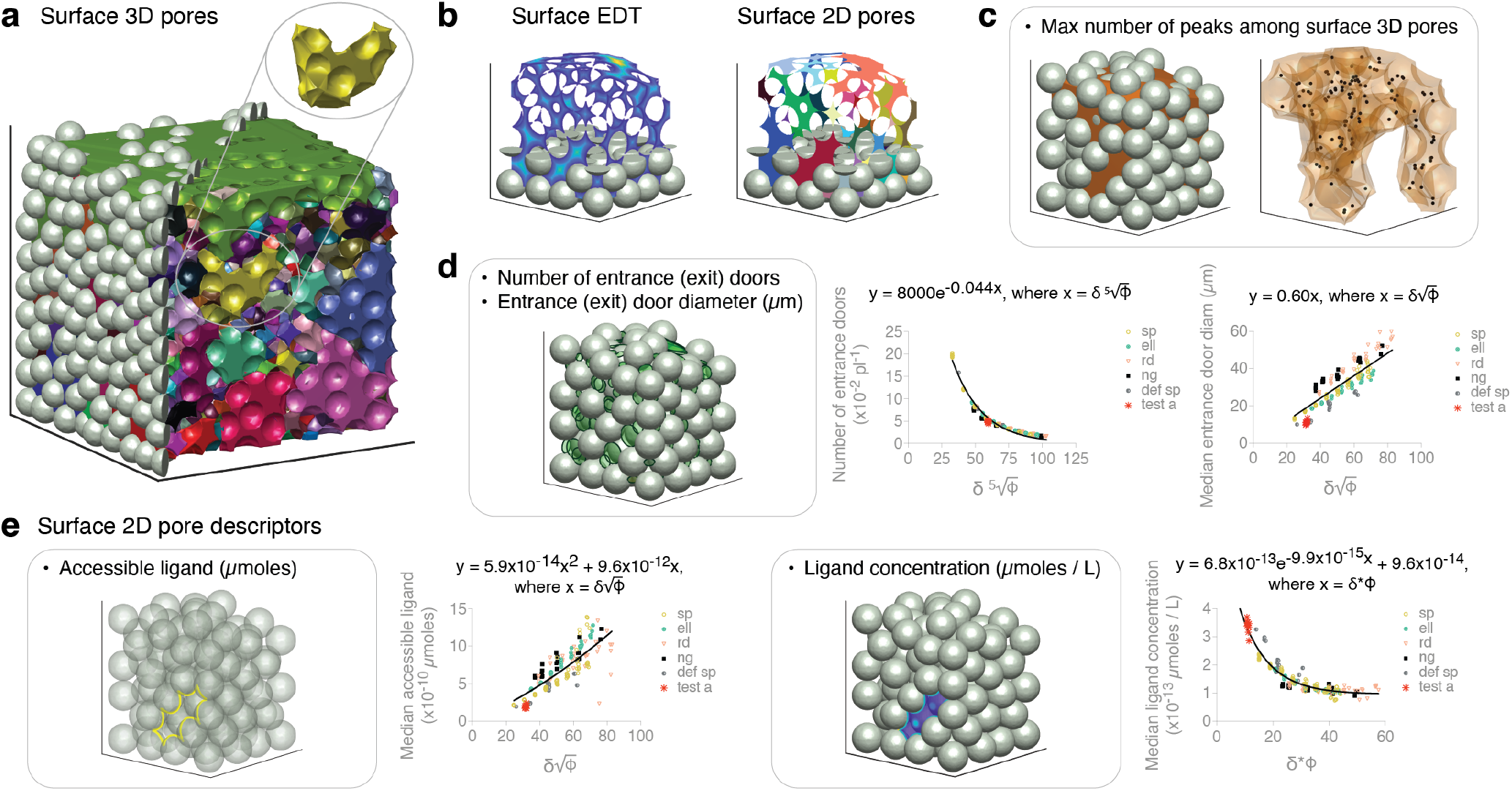
Surface 3D- and 2D-pore descriptors. **a**, Sample packed particle assembly (left, gray) with corresponding surface 3D pores (distinct colors), highlighting a yellow surface 3D pore. **b**, (left) Heatmap representation of the Euclidean distance transform (EDT) computed on the surface (2-manifold) of the void space, where lighter colors indicate regions that are farther from particles. (right) Surface 2D pores (distinct colors), representing the location of distinct openings in the structure. **c**, Maximum number of peaks in any single pore of a scaffold. **d**, Number and diameter (µm) of entrance (exit) doors within a scaffold (entrance doors represented as green circles). Equations predict the total number of entrance doors and the median entrance door diameter for monodisperse packings given δ and ϕ. **e**, (left) Accessible ligand (µmoles) of surface 2D pores, where yellow indicates a trail of ligand along particle surfaces that outlines the 2D pore. Equation predicts the median accessible ligand of surface 2D pores for monodisperse packings given δ and ϕ. (right) Ligand concentration (µmoles / L) of surface 2D pores computed from the ligand heatmap described in Figure 1g; color key shown in Figure 1g iv. Equation predicts the median ligand concentration of surface 2D pores for monodisperse packings given δ and ϕ. δ, particle diameter; ϕ, void volume fraction; sp, spheres; ell, ellipsoids; rd, rods; ng, nuggets; def sp, deformable spheres; test a, unbiased estimator of softer deformable spheres with δ = 100, λ= 400, μ = 50. N = 10 domains (sp, def sp, test a) and N = 4 – 5 domains (ell, rd, ng).

Entrance (or exit) doors along the surface of the void space are centered at peaks along the 2D surface of the void space (Figure 3d), where such peaks are points on the surface that are equally far from three or more nearby particles(1). Unlike interior doors that represent the narrowest regions in space, entrance doors represent the widest regions and may overlap with one another. The number of entrance doors per picoliter is a predictable feature across scaffolds given a transformation on particle diameter and void volume fraction (Figure 3d):

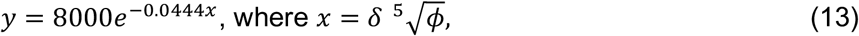

where *δ* and *ϕ* are particle diameter and void volume fraction, respectively (R^2^ = 0.9789, NRMSE = 0.0317). We also report a best fit line for predicting the median entrance door diameter across scaffolds:

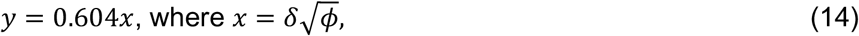

where *δ* and *ϕ* are as above (R^2^ = 0.7919, NRMSE = 0.528). Although median entrance door diameter trends upward for all packed particle groups, we begin to see that median surface pore information is more difficult to predict using particle size and void volume fraction alone.

Since surface 2D pores represent distinct openings into and out of the scaffold through which an infiltrating cell would pass, we report accessible ligand and ligand concentration through these openings to aid potential studies of cell adhesion patterns at the surface. Accessible ligand for surface 2D pores amounts to outlining each opening with ligand (Figure 3e, left), while ligand concentration of these openings is displayed like a window with overlayed ligand heatmap (Figure 3e, right). Resulting equations for predicting median accessible ligand (µmoles) and median ligand concentration (µmoles / L) along the surface are:

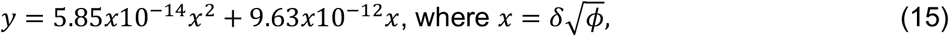

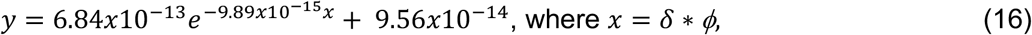

where *δ* and *ϕ* are as above. As with predicting median entrance door diameter, median accessible ligand is more difficult to predict across scaffold groups (R^2^ = 0.6836, NRMSE = 0.8078), although we see a better fit for median ligand concentration (R^2^ = 0.8276, NRMSE = 0.00429).

### Paths through void space

Flow, mass transport, and object migration are commonly studied in packed particle systems, which motivates our study of paths along medial axis curves of the void space. We define paths as the shortest routes from the center of the void space to the surface along 1D ridges (Figure 4a, b). Each path traverses a set of pores on its way out (Figure 4c). We report several path descriptors, including the total number of paths per picoliter (Supplementary Figure 4), path length, and two measurements of tortuosity (Figure 4d). Tortuosity-by-length is the simplest definition of tortuosity, 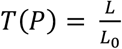, where *P* is a path, *L* is the length of the path, and *L*_0_ is the distance between the endpoints of the path. Tortuosity-by-volume is the volume of the convex hull containing the set of path points. Unlike a path-to-chord ratio, a volume-based approach for describing tortuosity will scale with path size (Supplementary Figure 5a).

**Fig. 4.**
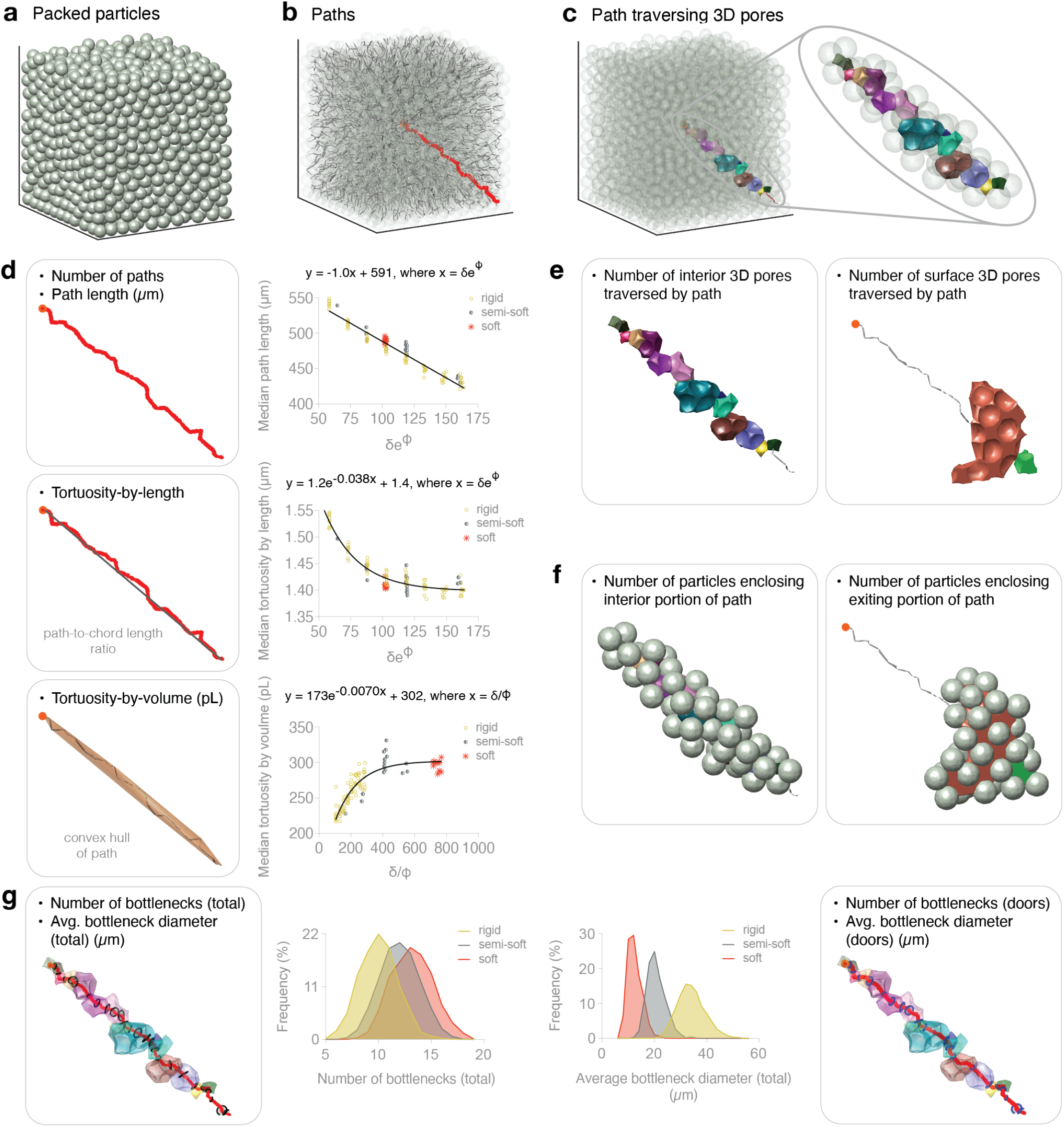
Path descriptors. **a**, Sample packed particle assembly. **b**, Paths starting at the center of the scaffold and traveling the shortest distance through pores to each exit point; highlighting a sample path in red. **c**, Highlighting 3D pores traversed by the sample path. **d**, Length (µm), tortuosity-by-length (i.e., arc-to-chord ratio), and tortuosity-by-volume (pL) (i.e., convex hull) of paths in scaffolds. Equations predict the median of these values given δ and ϕ for monodisperse spheres of increasing deformability. **e**, Number of interior and surface 3D pores traversed by paths in scaffolds. **f**, Number of particles enclosing the interior-pore and surface-pore portion of paths in scaffolds. **g**, Number and average diameter (µm) of bottlenecks (black circles) representing all narrow points, including doors, along paths in scaffolds; sample path includes transparent pores. Showing distribution plots for monodisperse spheres of increasing deformability. **h**, Number and average diameter (µm) of door-only bottlenecks (blue circles) along paths in scaffolds; sample path includes transparent pores. δ, particle diameter; ϕ, void volume fraction; semi-soft, λ= 1,000, μ = 250; soft, λ= 400, μ = 50. N = 10 domains (rigid, semi-soft, soft).

**Fig. 5.**
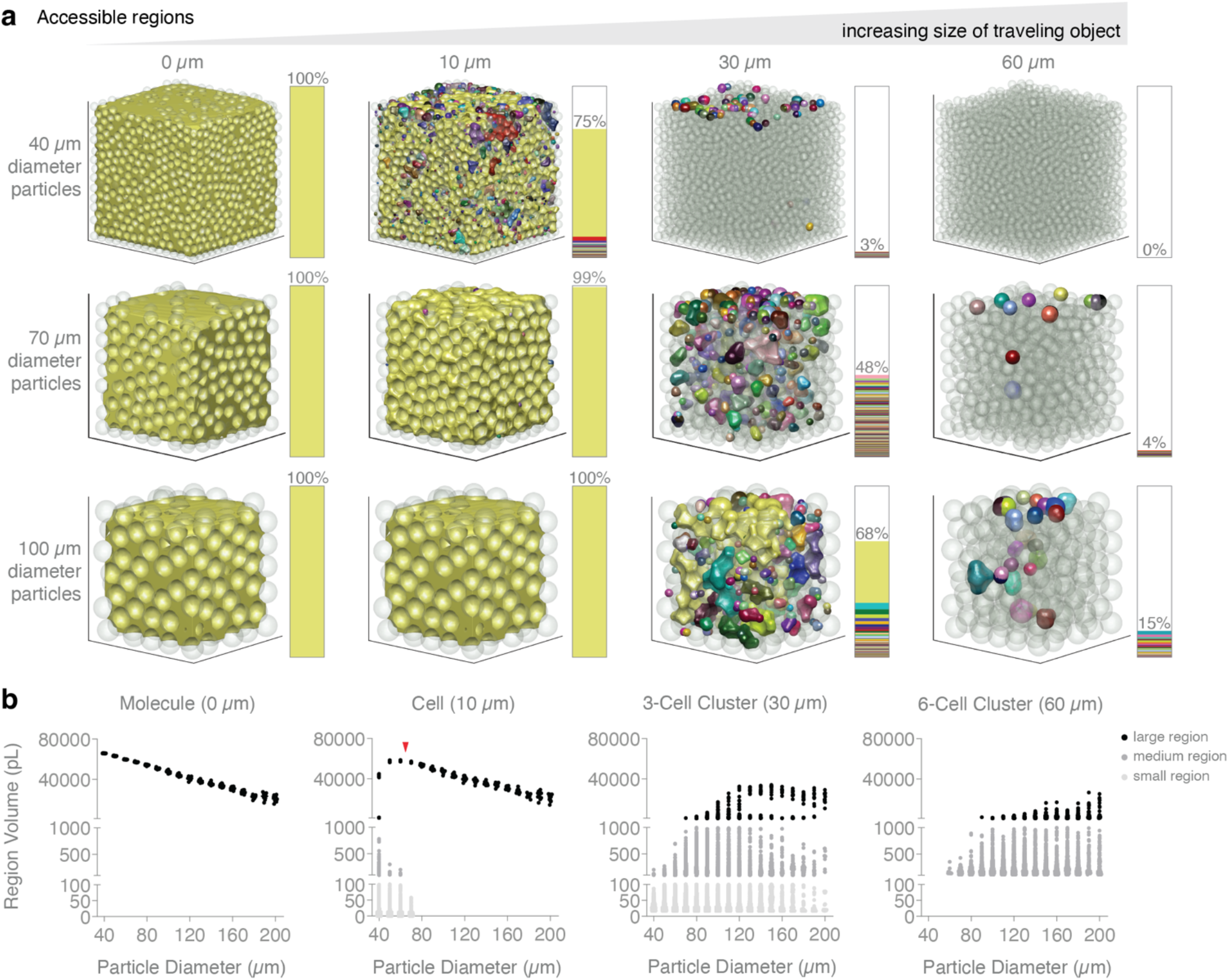
Accessible regions descriptors. **a**, Showing visualizations of accessible regions (distinct colors) for 0, 10, 30, and 60 µm diameter spherical objects traveling through the void space of packed particle assemblies comprising 40, 70, and 100 µm diameter spheres. Stacked bar chart illustrates the proportional volume of each region, with gray number indicating the cumulative proportion of accessible void space. **b**, Scatter plots of corresponding accessible region volumes categorized by size (small, medium, large). Plot titles include biological approximations to the size of the traveling object. Red arrow in Cell plot indicates particle diameter threshold after which accessible region volumes follow that of Molecule plot. N = 10 domains for each particle size.

An output comparison between the two metrics is presented in Supplementary Figure 5b. To study these descriptors, we consider packings of spherical particles with ranging deformability: rigid, semi-soft (Lamé parameters λ = 1,000, μ = 250), and soft (Lamé parameters λ = 400, μ = 50). Applying transformations to the independent variables produces the following regression models for path length (µm), tortuosity-by-length, and tortuosity-by-volume (pL), respectively:

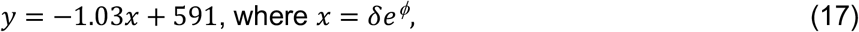

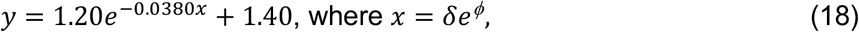

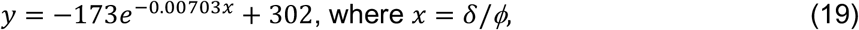

where *δ* is particle diameter and *ϕ* is void volume fraction (R^2^ is 0.9730, 0.9077, and 0.7993, respectively) (Figure 4d). To build additional intuition for paths, we provide visuals of paths and their corresponding convex hulls (tortuosity-by-volume) for packings comprising mixtures of rigid and soft spheres at different proportions (Supplementary Figure 6).

For each path, we report the number of interior pores and surface pores traversed by the path (Figure 4e), as well as the number of particles enclosing both the interior and exiting portion of the path (Figure 4f). Recall that 1D ridges are uniquely defined by their equidistant particles, and each 1D ridge contains a narrowest point along the curve. Since paths are formed by contiguous 1D ridges, we identify these narrowest points as bottlenecks, for which we report the total number and average diameter of bottlenecks per path (Figure 4g, left). Frequency distributions are displayed for rigid, semi-soft, and soft particle domains, indicating that softer particles produce increased numbers of bottlenecks per path with smaller bottleneck diameters, as expected (Figure 4g). A subset of these bottlenecks are doors, which typically exist between pores. Occasionally, a door may exist on the branch of a looped backbone within a single large pore. We report the total number and average diameter of doors per path (Figure 4, right).

### Size-dependent accessible regions

Finally, we consider how the size of an object impacts where it can move within the void space. We study four spherical object species of varying diameters: 0, 10, 30, and 60 µm. For each object, we identify all distinct regions of the void space in which the object can fit. Figure 5a provides a visual representation of these accessible regions for packings comprising 40, 70, and 100 µm diameter rigid particles. Distinct regions are highlighted by unique colors, and, by definition, an object cannot move between regions. As an object becomes larger, the available regions become increasingly disjoint and sparse. Beside each scaffold image, we provide a proportional bar chart of the regions, where the percentage label indicates the cumulative space available to the object.

For biological relevance, our object species are akin to molecules (~ 0 µm), cells (10 µm), 3-cell clusters (30 µm), and 6-cell clusters (60 µm). We study scaffolds comprising monodisperse rigid spheres ranging from 40 to 200 µm in diameter, and for each species category, we report the volumes of all regions, annotating them as small, medium, or large (Figure 5b). Since void space is defined by the convex hull of particle centers, the total void space volume decreases as particle size increases. This trend is evident in the 0 µm plot of Figure 5b, where we see a downward trajectory in the data. In a packed particle assembly, door diameter dictates local accessibility for a traveling sphere. Generally, a traveling object can move unimpeded if it can fit through the doors of the smallest pore type in the system. For monodisperse packings of spheres, the smallest pore type is formed by four touching spheres (Supplementary Figure 7a). We know the relationship between particle diameter, *d*_*particle*_, and door diameter, *d*_*door*_, is 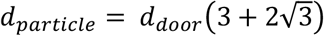 (Supplementary Figure 7b). Therefore, for a traveling spherical object of diameter *d*_*door*_ moving through a system of monodisperse packed spheres, we can predict the minimum particle diameter for which the object should be able to move freely throughout the void space. Such a scenario would generate a single accessible region that equals the total void space volume. For a 10 µm diameter object, the minimum particle diameter is approximately 65 µm, which corresponds to the tapering-off point (red arrow) in our 10 µm plot – the point after which datapoints mirror those in our 0 µm plot (Figure 5b). For 30 and 60 µm diameter objects, these diameter thresholds are approximately 194 and 388 µm, respectively.

## Discussion

We expand our initial study of porosity presented in Riley et al. 2023 by introducing a set of new metrics to evaluate the relationship between particle packing and void space. The first of these descriptors capture aspects of 3D pores – including size, shape, connectivity, and how particle surfaces are perceived from within pores. We introduce a method to quantify the vertices, edges, and faces of 3D pores, allowing their shapes to be approximated to simpler polyhedra. Our findings are validated for the most fundamental pore type that is known to approximate a tetrahedron (Figure 1f, inset). For rigid spheres, results reveal distributional patterns for pore shape descriptors (vertices, edges, faces; Figure 1f) and connectivity metrics (hallways, crawl spaces; Figure 2b) that generally follow linear combinatorial scaling laws grounded in graph theory.

Our metrics for studying phenomenon at particle-surface interfaces are reported in units of quantity and concentration, applied here to ligand protein distribution as an example relevant to biomaterials engineering. However, our approach provides a general framework that scales linearly with initial condition, which allows concepts to be broadly translatable. For example, understanding absolute quantities along particle surfaces from the perspective of local pores (Figure 1g, *ii*) is relevant to systems involving adsorption and filtration(38); ligand hotspots (Figure 1g, *vi*) highlight reaction centers in heterogeneous catalysis and capacitive systems(39); and ligand heatmaps (Figure 1g, *iii*) resemble a snapshot of a diffusion process, with relevance to heat and mass transfer in packed bed reactors(40).

We perform empirical modeling to derive predictive relationships between particle properties and pore descriptors. For the range of simulated, monodisperse domains in our study, transformations combining average particle diameter (or the diameter of an equal-volume sphere), *δ*, and void volume fraction, *ϕ*, capture enough structural information to produce well-fit regression models for the descriptors reported. Interestingly, different descriptors require different transformations to achieve strong predictive performance, yet the functional forms remain simple. The transformation 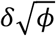 appears to be a reliable predictor for all reported interior descriptors that scale with length. While the regression equations presented herein are fit to data from monodisperse particle packings, we anticipate that incorporating a term to account for the standard deviation in particle diameter will yield generalizable models for polydisperse scaffolds, as demonstrated in Riley et al. 2023. However, analysis of polydisperse systems is beyond the scope of this study.

To extend our understanding of scaffold accessibility, we include three additional categories of metrics: openings into and out of the structure (i.e., surface descriptors), paths that run centrally through the porous network, and distinct void space regions that are accessible to objects of specified size. For surface metrics, our transformations yield the weakest regression performance, highlighting the variability introduced by container edge effects. Incorporating additional particle packing attributes – alongside particle diameter and void volume fraction – may lead to stronger predictive relationships.

Paths and accessible regions calculations offer practical insight into porosity, allowing packed particle engineers to select particle sizes that will optimize unrestricted movement or local accessibility.

We choose to focus our analyses on a subset of descriptors that are of particular interest; however, we leave the scientific community to explore undiscovered patterns in the large body of descriptor data. While our simulated dataset incorporates variations in particle size, shape, and stiffness, expanding the dataset could help refine or strengthen the conclusions presented here. Furthermore, our analysis describes static particle assemblies, but future work might explore how descriptor patterns evolve as particles undergo physical rearrangement.

Visuals of several descriptors for perfect packing are shown in Supplementary Figure 8. Additional straight-forward descriptors outputted by LOVAMAP are presented in Methods. A complete list of regression equations is provided in Supplementary Table 4.

## Methods

### Generating particle domains

Our packed particle assemblies are simulated using SideFx Houdini software, which we describe in detail in Riley et al. 2023(1). Briefly, particles are dropped into a 600 × 600 × 600 µm container and left to settle according to rigid body dynamics at default settings. We refer the reader to Riley et al. for details regarding generation of spheres, ellipsoids (prolate spheroids), nuggets, and rods (cylinders). Domains are discretized on a uniform Cartesian grid with mesh size of *dx* = 2, and data is converted to appropriate format for LOVAMAP input(1).

### Forming pores

Void space segmentation to generate 3D pores and surface 2D pores is described in detail in Riley et al. 2023(1). Briefly, we define the void space as the non-particle portion of the convex hull of particle centers. We compute the EDT on the void space for each particle to identify subtypes of the void-space medial axis that are used as a basis for segmentation. 2D ridges refer to medial axis points that are equidistant to two particles, 1D ridges are points that lie at intersections of 2D ridges, and peaks are points that lie at the intersection of 1D ridges. 1D ridges are equidistant to three or more particles, while peaks are equidistant to four or more particles and represent local center points in the void space. 1D ridges and peaks form a 3D graph of edges and nodes, respectively. 3D pores are first defined by their peaks, where multiple peaks are associated to the same pore if the distance between neighboring peaks (i.e., those that share a 1D ridge) is less than the sum of their EDT values. Pore boundaries are defined at the minimum EDT value along 1D ridges that are flanked by peaks not associated to the same pore. A nearest-neighbors algorithm is then applied to pore backbones to fill the volume and finalize interior 3D pores. To generate surface 2D pores, we compute the EDT along the “2D surface” of the void space to identify the surface medial axis and subsequent surface peaks, which are local maximums of the EDT. We next apply a threshold cutoff to form surface-backbone “islands” that define surface 2D pores. The completed 2D pores are then formed using a nearest-neighbors algorithm.

Surface 3D pores are finalized by combining interior 3D pores that extend to the same surface 2D pore.

### Computing descriptors

#### Global descriptors

The following global descriptors are formally defined in Riley at al. 2023(1): *Number of particles, void volume fraction*, and *number of 3D pores (surface + interior)*. Below we include all additional global descriptors outputted by LOVAMAP.

##### Void volume fraction of interior pores

The cumulative volume of interior pores divided by the total volume of the convex hull of particle centroids.

##### Void area fraction

30 evenly-spaced 2D z-slices are sampled within the middle two-thirds of the scaffold along the z-axis. A z-slice of the domain matrix is all rows (x) and all columns (y) along the specified page (z), and for each z-slice, we compute the ratio of void voxels to all voxels within the convex hull. The final void area fraction reports the mean across all z-slices. The number of z-slices is an adjustable parameter.

##### Particle packing fraction

The cumulative volume of particle voxels contained within the convex hull of particle centroids divided by the total volume of the convex hull, i.e., 1 – void volume fraction.

##### Number of interior 3D pores

We report the total number of interior 3D pores within the packed particle domain; this excludes surface 3D pores.

##### Number of interior 3D pores / Number of particles surrounding interior pores

The number of interior 3D pores divided by the number of unique particles that surround all interior 3D pores.

##### Maximum number of equidistant particles

For a given scaffold, we search for the peak that is equidistant to the greatest number of particles, and we report the particle count.

##### Number of particle contacts

Particle contacts are defined as particles that are within distance *dx* (i.e., one voxel) from one another. We report the total number of particle contacts in a scaffold.

##### Number of interior doors

We report the total number of interior doors, where an interior door is a circle centered along a 1D ridge that lies between neighboring pores and extends to equidistant particles.

##### Number of paths

We report the total number of paths in a scaffold, where a path follows 1D ridges from the center of the scaffold along the shortest distance to an endpoint on the surface of the void space.

##### Particle adjacency matrix maximum eigenvalue

The particle adjacency matrix is an *N x N* square Boolean matrix that is constructed using 2D-ridge data, where *N* is the number of particles and a value of *true* is stored at (*i, j*) when particles *i* and *j* are neighbors that share a 2D ridge. We report the maximum eigenvalue of this matrix, which is bounded below by the average number of edges (2D ridges) per node (particle) and bounded above by the maximum number of edges for a single node.

##### Peak adjacency matrix maximum eigenvalue

The peak adjacency matrix is an *N x N* square Boolean matrix, where *N* is the number of peaks and a value of *true* is stored at (*i, j*) when peaks *i* and *j* reside in the same pore. We report the maximum eigenvalue of this matrix, which is bounded below by the average number of peaks contained within a single pore and bounded above by the maximum number of peaks associated with a single pore.

##### Pore adjacency matrix maximum eigenvalue

The pore adjacency matrix is an *N x N* square Boolean matrix, where *N* is the number of pores and a value of *true* is stored at (*i, j*) when pores *i* and *j* are neighbors that share a 1D ridge (hallway, door). We report the maximum eigenvalue of this matrix, which is bounded below by the average number of pore neighbors and bounded above by the maximum number of neighbors for a single pore in the scaffold.

##### Number of entrance (exit) doors

We report the total number of entrance (or exit) doors, which is equivalent to the total number of peaks along the 2D surface of the void space.

##### Maximum number of peaks among surface 3D pores

Particle composition in relation to edge effects of the holding container drastically influences the landscape of surface pores (Supplementary Figure 3). To capture an element of this phenomenon, we look at the number of peaks contained within surface 3D pores, since the number of peaks scales with the size and complexity of pores. For each domain, we report the peak count of the surface 3D pore that contains the most number of peaks.

##### Ligand hotspots volume fraction

Our scaffold is converted into a ligand heatmap by applying an averaging-filter over our ligand data using a spherical kernel whose diameter is 1/60^th^ the x-axis length of the scaffold (10 µm in our experiments). Hotspots are defined as void space voxels in the ligand heatmap containing ≥ 2,167,200 ligand molecules, which corresponds to 90% of the maximum ligand count across all heatmaps. To obtain the ligand hotspots volume fraction, we divide the number of hotspot voxels by the total number of voxels in the void space. Ligand heatmaps are a function of the domain mesh size (dx = 2) and ligand concentration (500 µmoles / L), both of which are held constant in our simulated runs. Hotspot threshold, kernel size, and ligand concentration are adjustable parameters.

#### 3D pore descriptors

The following 3D pore descriptors are formally defined in Riley at al. 2023(1): *Volume (pL), longest length (µm), average internal diameter (µm), aspect ratio, largest enclosed sphere diameter (µm), number of surrounding particles, length of PCA ellipsoid axis 1, 2, and 3*, and *isotropy (anisotropy)*. Below we include additional 3D pore descriptors outputted by LOVAMAP.

#### Surface area (µm^2^ / 1000)

The surface voxels of a pore are the single-voxel-thick outer layer of the pore. We count the surface voxels of the pore and multiply by dx^2^ as a crude approximation to surface area. Division by 1000 allows for comparison to volume, which is reported in pL.

#### Characteristic length (µm)

Volume (µm^3^) / Surface area (µm^2^). For reference, the characteristic length of a sphere with radius r is r/3, while a cube with height h is h/6.

#### Number of peaks

We count the number of peaks within a given pore.

#### Particle surface area (µm^2^)

The surface voxels of a pore are the single-voxel-thick outer layer of the pore. We count the surface voxels that neighbor a particle and multiply by dx^2^.

#### Open (non-particle) surface area (µm^2^)

The surface voxels of a pore are the single-voxel-thick outer layer of the pore. We count the surface voxels that do not neighbor a particle and multiply by dx^2^.

#### Mean local thickness

Briefly, the local thickness of a point in a pore is equal to the diameter of the largest 2D-ridge-centered sphere that encloses it, and we arrive at mean local thickness by integrating and normalizing by volume. More specifically, we first aim to construct a thickness matrix per pore that assigns a ‘thickness’ value to each voxel in the pore. To obtain thickness values for a given pore, we start by identifying all 2D-ridge voxels that are contained within the boundary of the pore. The thickness matrix at these 2D-ridge voxels are set to the voxels’ EDT value multiplied by 2, and the remaining voxels are initialized to zero. For each 2D-ridge voxel, *v*_*i*_, within the pore, we compute the distance between *v*_*i*_ and all other pore voxels in order to identify the pore voxels, {*s*_*i*_}, that lie within a sphere centered at *v*_*i*_ with radius equal to the EDT value at *v*_*i*_. The thickness values for voxels in {*s*_*i*_} are updated to equal the maximum between their current value and the EDT of *v*_*i*_ multiplied by 2 (i.e., diameter of the sphere). The mean local thickness is the integral of local thickness over the pore divided by the volume of the pore. Our methods are adapted from Hildebrand T and Ruegsegger P, 1997(33).

##### Number of edges

We count the number of edges of the pore, where edges are formed at the intersection of particle-faces and non-particle-faces. This amounts to counting the cumulative number of particles associated with each hallway (1D ridge) exiting the pore.

##### Number of vertices (same as ‘crawl spaces’)

We count the number of particle-pair neighbors surrounding the pore, which corresponds to the number of 2D ridges exiting the pore.

##### Number of faces

We count the number of faces of the pore, where faces are formed by either surrounding particles or doors. This amounts to summing the number of surrounding particles plus the number of hallways (or doors).

##### Particle surface area / open surface area

Particle surface area (µm^2^) / Open (non-particle) surface area (µm^2^)

##### Ellipsoid classification

For each pore, we perform PCA by computing the covariance matrix from the pore’s voxel coordinates to identify its three orthogonal eigenvectors and eigenvalues. These principal axes are then used to classify the pore shape according to the following key: 1 = sphere, assigned when axis 3 is at least 80% the length of axis 1 (i.e., all axes are similar in length); 2 = pancake, assigned when axis 1 and 2 are similar relative to axis 3, and axis 3 is less than 1/3 the length of axis 1; 3 = tube, assigned when axis 2 and 3 are similar relative to axis 1, and axis 2 is less than 1/3 the length of axis 1; 0 = undefined, assigned when none of the above criteria are met.

##### Number of hallways

We count the number of 1D ridges that exit the pore, which is equivalent to the number of doors that separate a pore from its neighboring pores.

##### Number of crawl spaces (same as ‘vertices’)

We count the number of 2D ridges that exit the pore, which corresponds to the number of particle-pair neighbors surrounding the pore.

##### Number of connected 3D pores

We report the number of neighboring 3D pores that share a 1D ridge with the pore, i.e., are accessible through a hallway (or door).

##### Number of surrounding 3D pores

We report the number of neighboring 3D pores that share a 2D ridge with the pore, i.e., are accessible through a crawl space.

##### Normalized neighbors

Number of hallways / Number of crawl spaces

##### Largest door diameter (µm)

Diameter of the largest door associated with the pore, where a door is a circle centered along a hallway (1D ridge) that lies at the boundary of the pore and extends to equidistant particles.

##### Smallest door diameter (µm)

Diameter of the smallest door associated with the pore, where a door is a circle centered along a hallway (1D ridge) that lies at the boundary of the pore and extends to equidistant particles.

##### Ligand concentration (µmoles / L)

Our scaffold is converted into a ligand heatmap by applying an averaging-filter over our ligand data using a spherical kernel whose diameter is 1/60^th^ the x-axis length of the scaffold. Ligand concentration here represents the average ligand concentration within a pore computed from the ligand heatmap. For a given pore, we sum the values of the ligand heatmap that correspond to the voxels of the pore, then divide by the volume of the pore. We convert to µmoles / L using 1 µm^3^ = 10^−15^ L and 1 µmole = 6.02 × 10^17^ molecules. Kernel size is an adjustable parameter.

##### Accessible ligand (µmoles)

For a given pore, accessible ligand represents the amount of ligand on particle surfaces that is accessible from inside of the pore. We first locate the particle voxels that enclose the pore. The total count of these voxels is then converted to a surface area, which is then multiplied by a thickness representing how far into a particle a cell can reach. This volume is then converted into µmoles of ligand using an experimentally-derived particle ligand concentration of 500 µmoles / L and a conversion of 1 µm^3^ = 10^−15^ L. Particle ligand concentration is an adjustable parameter.

##### x-, y-, z-centroid coordinate

The (x, y, z) coordinate of the pore’s centroid.

#### Other descriptors

##### Particle diameter (µm)

We report the diameter of a sphere with the same volume as the particle.

##### Coordination number: neighboring particles

For a given particle, this descriptor returns the number of surrounding particles that share a 2D ridge with the particle.

##### Coordination number: touching particles

For a given particle, this descriptor returns the number of surrounding particles that are within a distance *dx* (i.e., one voxel) from the particle.

##### Entrance (Exit) door diameter (µm)

An entrance (or exit) door is represented by a circle at an opening into and out of the scaffold. It is centered at peaks along the “2D surface” of the void space and extends to equidistant particles. We report the diameter of the door.

##### Interior door diameter (µm)

An interior door is represented by a circle at the boundary between neighboring pores. It is centered at the voxel with the minimum EDT value along the hallway (1D ridge), and it extends to equidistant particles. We report the diameter of the door.

##### Path length (µm)

A path follows 1D ridges from the center of the scaffold along the shortest distance to an endpoint on the surface of the void space. We sum the lengths of the 1D-ridge segments that make up the path. The arc length of each 1D ridge is computed by approximating 1D ridges using least squares splines.

##### Tortuosity-by-length

For each path, we report the classic arc-to-chord ratio. The arc length is equivalent to our computation of path length; the chord length is the Euclidean distance between the start of the path (the central peak of the scaffold) and the end of the path at the surface of the void space.

##### Tortuosity-by-volume (pL)

For each path, we use MATLAB methods to compute the volume of the 3D convex hull of all path voxels.

##### Number of interior 3D pores traversed by path

We identify the unique interior 3D pores associated with the 1D-ridge segments that form the path, then report the total number of interior pores.

##### Number of surface 3D pores traversed by path

We identify the unique surface 3D pores associated with the 1D-ridge segments that form the path, then report the total number of surface 3D pores.

##### Number of particles enclosing interior portion of path

We report the number of particles surrounding interior 3D pores traversed by the path; this excludes particles surrounding surface 3D pores traversed by the path.

##### Number of particles enclosing exiting portion of path

We report the number of particles surrounding surface 3D pores traversed by the path; this excludes particles surrounding interior 3D pores traversed by the path.

##### Number of bottlenecks (total)

We define bottlenecks as the narrowest point along each 1D-ridge segment of a path, which corresponds to the point with the smallest EDT value. For this descriptor, we report the total number of all bottlenecks.

##### Average bottleneck diameter (total) (µm)

We report the average bottleneck diameter for a given path, where bottlenecks are the narrowest point along each 1D-ridge segment of the path.

##### Number of bottlenecks (doors)

Doors are narrowed points along 1D ridges that span neighboring pores, making them a specific subset of bottlenecks. We report the number of doors along a path. On occasion, a door may exist within a 3D pore if it lies along a 1D ridge that is flanked by two peaks within the pore that satisfy the door criteria (i.e., the sum of the peaks’ EDT values is less than the distance between them). In these scenarios, there will be another route along the pore’s backbone for which the door criteria is not satisfied such that the pore still captures an open space.

##### Average bottleneck diameter (doors) (µm)

We report the average diameter of doors along a given path.

##### Surface ligand concentration (µmoles / µm^2^)

Our scaffold is converted into a ligand heatmap by applying an averaging-filter over our ligand data using a spherical kernel whose diameter is 1/60^th^ the x-axis length of the scaffold. For each surface 2D pore, we sum the values in the ligand heatmap that correspond to voxels of the 2D pore and convert to micromoles using a conversion of 1 µmole = 6.02 × 10^17^ molecules. We then divide by the surface area of the pore. Kernel size is an adjustable parameter.

##### Surface accessible ligand (µmoles)

This descriptor represents the amount of ligand that is accessible to an infiltrating object along the 2D surface of each scaffold opening. For each surface 2D pore, we locate the particle voxels that surround the pore by checking which pore voxels neighbor a particle voxel. The total count of these voxels is then converted to a surface area, which is then multiplied by the user-inputted value representing how far into a particle a cell can reach (shell thickness). This volume is then converted into µmoles of ligand using an experimentally-derived particle ligand concentration of 500 µmoles / L and a conversion of 1 µm^3^ = 10^−15^ L. Particle ligand concentration is an adjustable parameter.

##### Available regions data (pL)

The goal is to isolate all distinct regions of the void space through which an object of a specified size can fit. We select four spherical object ‘species’ with the following diameters: 0 µm, 10 µm, 30 µm, and 60 µm. 0 µm objects are taken as the size-scale of molecules, while 10 µm objects are akin to cells, 30 µm objects to 3-cell clusters, and 60 µm objects to 6-cell clusters. For each species size category, we locate the 1-manifold void space backbone voxels associated with an EDT value that is greater than or equal to the radius of the object. We then use a connected components approach to isolate unique islands of backbone clusters. For the smallest category, the EDT threshold is set to 0, and we locate the largest island to define the main backbone of the scaffold. All voxels not connected to this main backbone are not considered in analysis for any species category.

Once we identify the backbone of distinct regions through which the species can fit, we must gather the remaining void space voxels associated with each region. For each region, we scan each voxel, *v*_*b*_, of the backbone and gather the void space voxels whose distance to *v*_*b*_ is less than the EDT value at *v*_*b*_. This cumulative collection of void space voxels forms the complete regions. Finally, we compute the volume of each region.

## Statistical analysis

For fitting data to curves, we use a least-squares approach and report R^2^. Model performance is evaluated using the normalized root mean squared error (NRMSE) on a test group used as an unbiased measure of model generalization.

## Acknowledgements

Thank you to Katia Segura-Price for her help in generating scatter plots of our accessible regions data. We thank the National Institutes of Health, the National Institutes of Neurological Disorders and Stroke (1R01NS112940 (T.S.), 1R01NS079691 (T.S.), R01NS094599 (T.S.)) and the National Institute of Allergy and Infectious Disease (1R01AI152568 (T.S.)).

## Author contributions

L.R. headed the work by conceiving descriptors, writing the code, generating and analyzing data, writing the paper, and creating figures. E.L. processed, organized, and analyzed data, contributed to writing, and participated in discussion. P.C. provided computational expertise, optimized code for descriptors, designed and simulated input domains, and contributed to editing. D.A.A. processed data and contributed to writing and editing. T.S. guided the conception and development of LOVAMAP and descriptors, as well as provided input during manuscript preparation.

