## Supplementary Information for "Metrics for studying the porous void space of packed particles"

### Supplementary Tables

| Global | Definition |
|---|---|
| **General** | |
| Number of particles | Number of **particles** |
| Void volume fraction | Ratio of **void space volume** over **total scaffold volume** |
| Void volume fraction of interior pores | Ratio of **total interior pore volume** over **total scaffold volume** |
| Average void area fraction | Average **void area fraction** across a sample of 2D-slices |
| Particle packing fraction | Ratio of **particle volume** over **total scaffold volume** |
| Number of 3D pores: surface + interior | Number of pores (**surface + interior**) |
| Number of 3D pores: interior only | Number of **interior pores** |
| Number of interior 3D pores / Number of particles surrounding interior pores | Ratio of **number of interior pores** over **number of particles bounding interior pores** |
| Max number of equidistant particles | **Maximum number of particles** equidistant to any single peak |
| **Internal Connectivity** | |
| Number of particle contacts | Number of **particle-particle contact points** |
| Number of interior doors | Number of **interior doors** |
| Number of paths | Number of **paths** that travel from the center of the scaffold to each exit door |
| Particle adjacency matrix eigenvalue | Eigenvalue of the **particle adjacency matrix** |
| Peak adjacency matrix eigenvalue | Eigenvalue of the **peak adjacency matrix** |
| 3D pore adjacency matrix eigenvalue | Eigenvalue of the **pore adjacency matrix** |
| **External Connectivity** | |
| Number of entrance (exit) doors | Number of **doors to enter or exit** the scaffold |
| Max number of peaks among surface 3D pores | **Maximum number of peaks** contained within a single surface 3D pore |
| **Ligand availability** | |
| Ligand hotspots volume fraction | Ratio of **ligand heatmap hotspots** over **total scaffold volume** |

**Supplementary Table 1 | LOVAMAP global descriptors.**

| 3D pore | Definition |
| --- | --- |
| | *Each measurement for the subsequent descriptors refers to a specific 3D pore* |
| **Size** | |
| Volume (pL) | **Volume** of the pore |
| Surface area ($\mu m^2$ / 1000) | **Surface area** of the pore |
| Characteristic length (µm) | Ratio of **volume** over **surface area** of the pore |
| Longest length (µm) | **Longest chord length** that spans two surface points of the pore |
| Average internal diameter (µm) | **Average width** along the internal backbone of the pore |
| Aspect ratio (longest length / avg. internal diam) | Ratio of **longest length** over **average internal diameter** |
| Largest enclosed sphere diameter (µm) | **Diameter of the largest sphere** that can fit entirely within the pore |
| Number of surrounding particles | Number of **particles** that enclose the pore |
| Number of peaks | Number of **peaks** contained within the pore |
| Particle surface area ($\mu m^2$) | **Surface area** of the pore's **physical boundary** formed by surrounding particles |
| Open (non-particle) surface area ($\mu m^2$) | **Surface area** of the pore's **conceptual boundary** in the void space |
| **Shape** | |
| Mean local thickness (µm) | The average **'thickness' measurement** computed at every point within the pore |
| Number of vertices (same as 'crawl spaces') | Number of **pore vertices** formed at particle contact points surrounding the pore |
| Number of edges | Number of **pore edges** formed at the intersection of the pore's physical (particle) and conceptual (non-particle) boundaries |
| Number of faces | Number of **pore faces** formed by surrounding particles and doors |
| Particle surface area / open surface area | Ratio of **particle-boundary surface area** over **non-particle-boundary surface area** |
| Ellipsoid axis 1 length (µm) | Length of the **largest axis** of the PCA ellipsoid computed from the pore's point cloud |
| Ellipsoid axis 2 length (µm) | Length of the **second largest axis** of the PCA ellipsoid computed from the pore's point cloud |
| Ellipsoid axis 3 length (µm) | Length of the **third largest axis (Axis 1)** of the PCA ellipsoid computed from the pore's point cloud |
| Ellipsoid classification | 0 = undefined shape, 1 = spherical shape, 2 = pancake-like shape, 3 = tube-like shape |
| **Connectivity** | |
| Number of hallways | Number of **exits through doors** of the pore |
| Number of crawl spaces (same as 'vertices') | Number of **exits through touching particles** surrounding the pore |
| Number of connected 3D pores | Number of **neighboring pores accessible through the doors** of the pore |
| Number of surrounding 3D pores | Number of **neighboring pores accessible through the doors and crawl spaces** of the pore |
| Normalized Neighbors (# hallways / # crawl spaces) | Ratio of the **number of hallways** over the **number of crawl spaces** |
| Largest door diameter (µm) | Diameter of the **largest door** of the pore |
| Smallest door diameter (µm) | Diameter of the **smallest door** of the pore |
| **Ligand availability** | |
| Ligand concentration (µmoles / L) | **Effective ligand concentration** of the pore as sensed by a migrating object |
| Accessible ligand (µmoles) | **Cumulative amount of ligand** on particle surfaces that bound the pore |
| **Directionality / Location** | |
| Isotropy (Anisotropy) | Value between 0 and 1, where **0 indicates** that the pore is oriented **perpendicularly to the average** orientation across all pores and **1 indicates** that the pore is orientated **in the same direction as the average** pore orientation |
| x-, y-, z-centroid coordinate | (x,y,z) coordinate of the **centroid** of the pore |

**Supplementary Table 2 | LOVAMAP 3D pore descriptors.**

| Other | Definition |
|---|---|
| Particles | *Each measurement for the subsequent descriptors refers to a specific particle* |
| Particle diameter (µm) | **Diameter** of the particle |
| Coordination number: neighboring particles | Number of particles that are **neighboring (close or touching)** the particle |
| Coordination number: touching particles | Number of particles that are in **contact with (touching)** the particle |
| Connectivity | *Each measurement for the subsequent descriptor refers to a specific door along the surface of the void space* |
| Entrance (Exit) door diameter (µm) | Diameter of the **entrance (exit) door** at the surface of the scaffold |
| | *Each measurement for the subsequent descriptor refers to a specific door within the scaffold* |
| Interior door diameter (µm) | Diameter of the **interior door** within the scaffold |
| | *Each measurement for the subsequent descriptor refers to a specific crawl space within the scaffold* |
| Crawl space width (µm) | Width of the **crawl space** within the scaffold |
| Paths | *Each measurement for the subsequent descriptors refers to a specific path* |
| Path length (µm) | **Length** of the path, which travels from the center of the scaffold to an exit door |
| Tortuosity-by-length | Ratio of **path length** over **straight-line distance between start and end points** of the path |
| Tortuosity-by-volume (pL) | Volume of the **convex hull** of all points along the path |
| Number of interior 3D pores traversed by path | Number of **interior pores** that are traversed by the path |
| Number of surface 3D pores traversed by path | Number of **surface pores** that are traversed by the path |
| Number of particles enclosing interior portion of path | Number of **particles enclosing the interior pores** along the path |
| Number of particles enclosing exiting portion of path | Number of **particles enclosing the surface pores** along the path |
| Number of bottlenecks (total) | Number of **narrow regions** along the path (including doors) |
| Average bottleneck diameter (total) (µm) | **Average diameter of the narrow regions** along the path |
| Number of bottlenecks (doors) | Number of **doors** that are traversed along the path |
| Average bottleneck diameter (doors) (µm) | **Average diameter of doors** along the path |
| Surface 2D pores | *Each measurement for the subsequent descriptors refers to a specific 2D pore along the surface of the void space* |
| Ligand concentration (µmoles / L) | **Effective ligand concentration** of the surface 2D pore as felt by a migrating object |
| Accessible ligand (µmoles) | **Cumulative amount of ligand** on particle surfaces that bound the surface 2D pore |
| Accessible Regions | *Each measurement for the subsequent descriptors refers to a distinct region of void space* |
| 0 µm object region volumes (pL) | Volume of the **total void space** within the scaffold |
| 10 µm object region volumes (pL) | Volume of a region of void space throughout which a **10 µm spherical object** can move unimpeded |
| 30 µm object region volumes (pL) | Volume of a region of void space throughout which a **30 µm spherical object** can move unimpeded |
| 60 µm object region volumes (pL) | Volume of a region of void space throughout which a **60 µm spherical object** can move unimpeded |

**Supplementary Table 3 | LOVAMAP other descriptors.**

| Descriptor | $y$ Predicts Median Value of Descriptor | Equation | Transformation* | Function Form | $R^2$ | NRMSE | Source |
|---|---|---|---|---|---|---|---|
| **3D Pores** | | | | | | | |
| Number of 3D pores ($pL^{-1}$) | | $y = 0.405e^{-0.0452x}$ | $x = (\delta + \sigma)e^{\phi}$ | exponential decay | 0.9827 | 0.0715 | Riley et al. 2023 |
| Volume (pL) | ✓ | $y = 0.0234x^2 - 0.595x$ | $x = (\delta + \sqrt{\sigma})\sqrt{\phi}$ | $2^{nd}$ order polynomial | 0.9402 | 0.178 | Riley et al. 2023 |
| Surface area ($\mu m^2$ / 1000) | ✓ | $y = 0.00356x^2 - 0.0345x$ | $x = \delta\sqrt{\phi}$ | $2^{nd}$ order polynomial | 0.9381 | 0.0355 | Equation 1 |
| Longest length (μm) | ✓ | $y = 1.32x$ | $x = \delta\sqrt{\phi}$ | linear (increasing) | 0.9492 | 0.0045 | Equation 2 |
| Average internal diameter (μm) | ✓ | $y = 0.473x$ | $x = \delta\sqrt{\phi}$ | linear (increasing) | 0.9862 | 0.0033 | Equation 3 |
| Largest enclosed sphere (μm) | ✓ | $y = 0.490x$ | $x = \delta\sqrt{\phi}$ | linear (increasing) | 0.9802 | 0.0080 | Equation 4 |
| Mean local thickness (μm) | ✓ | $y = 0.307x$ | $x = \delta\sqrt{\phi}$ | linear (increasing) | 0.9687 | 0.1037 | Equation 5 |
| Particle surface area ($\mu m^2$) | ✓ | $y = 2.32x^2$ | $x = \delta\sqrt{\phi}$ | $2^{nd}$ order polynomial | 0.9113 | 0.0405 | Equation S1 |
| Open surface area ($\mu m^2$) | ✓ | $y = 1.43x^2 - 59.7x + 812$ | $x = \delta\sqrt{\phi}$ | $2^{nd}$ order polynomial | 0.956 | 8.24 | Equation S2 |
| Accessible ligand (μmoles) | ✓ | $y = 4.64x10^{-12}x^2$ | $x = \delta\sqrt{\phi}$ | $2^{nd}$ order polynomial | 0.9113 | 0.0405 | Equation 6 |
| Ligand concentration (μmoles / L) | ✓ | $y = 197e^{-0.0676x} + 34.2$ | $x = \delta\phi$ | exponential decay | 0.973 | 0.000973 | Equation 7 |
| Largest door diameter (μm) | ✓ | $y = 0.448x$ | $x = \delta\sqrt{\phi}$ | linear (increasing) | 0.9650 | 0.0289 | Equation 10 |
| Smallest door diameter (μm) | ✓ | $y = 0.245x$ | $x = \delta\sqrt{\phi}$ | linear (increasing) | 0.7177 | 0.0253 | Equation 11 |
| **Connectivity** | | | | | | | |
| Number of interior doors ($pL^{-1}$) | | $y = 1.14e^{-0.0423x}$ | $x = \delta e^{\phi}$ | exponential decay | 0.9888 | 0.0462 | Equation 8 |
| Interior door diameter (μm) | ✓ | $y = 0.364x$ | $x = \delta\sqrt{\phi}$ | linear (increasing) | 0.9461 | 0.0870 | Equation 9 |
| Number of particle contacts ($pL^{-1}$) | | $y = 0.799e^{-0.0417x}$ | $x = \delta e^{\phi}$ | exponential decay | 0.9869 | 0.0203 | Equation S3 |
| 3D pore adjacency matrix max eigenvalue | ✓ | $y = 0.000173e^{-0.00704x} + 0.0000235$ | $x = \delta/\phi$ | exponential decay | 0.7487 | 0.0678 | Equation 12 |
| Number of entrance doors ($pL^{-1}$) | | $y = 8000e^{-0.0444x}$ | $x = \delta^5\sqrt{\phi}$ | exponential decay | 0.9789 | 0.0317 | Equation 13 |
| Entrance door diameter (μm) | ✓ | $y = 0.604x$ | $x = \delta\sqrt{\phi}$ | linear (increasing) | 0.7919 | 0.528 | Equation 14 |
| Surface accessible ligand (μmoles) | ✓ | $y = 5.85x10^{-14}x^2 + 9.63x10^{-12}x$ | $x = \delta\sqrt{\phi}$ | $2^{nd}$ order polynomial | 0.6836 | 0.808 | Equation 15 |
| Surface ligand concentration (μmoles / L) | ✓ | $y = 6.84x10^{-13}e^{-9.89E-15x} + 9.56x10^{-14}$ | $x = \delta\phi$ | exponential decay | 0.8276 | 0.00429 | Equation 16 |
| **Paths** | | | | | | | |
| Number of paths ($pL^{-1}$) | | $y = 0.0420e^{-0.0251x}$ | $x = \delta e^{\phi}$ | exponential decay | 0.9788 | 0.157 | Equation S4 |
| Path length (μm) | ✓ | $y = -1.02x + 591$ | $x = \delta e^{\phi}$ | linear (decreasing) | 0.9730 | | Equation 17 |
| Tortuosity-by-length | ✓ | $y = 1.20e^{-0.03798x} + 1.398$ | $x = \delta e^{\phi}$ | exponential decay | 0.9077 | | Equation 18 |
| Tortuosity-by-volume (pL) | ✓ | $y = -173e^{-0.00703x} + 302$ | $x = \delta/\phi$ | exponential decay | 0.7993 | | Equation 19 |

* $\delta$ = average particle diameter (or diameter of sphere with equal volume)
$\phi$ = void volume fraction of particle packing
$\sigma$ = standard deviation of particle diameter

### Supplementary Table 4 | Regression equations.

### Supplementary Figures

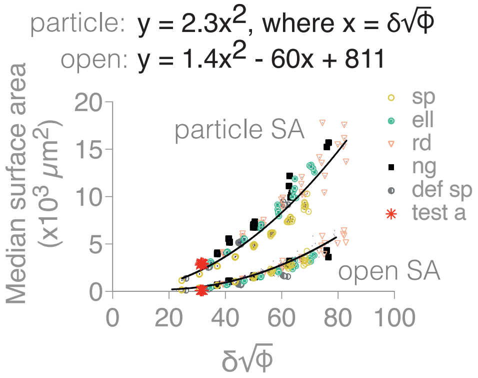


**Supplementary Figure 1 | Particle surface area and open surface area of pores.** Empirically-derived quadratic equations for predicting median particle-based surface area ($R^2$ = 0.9113) versus open surface area ($R^2$ = 0.9560) as a function of the transformation, respectively:

$$y = 2.32x^2, \text{ where } x = \delta\sqrt{\phi}, \tag{S1}$$

$$y = 1.43x^2 - 59.7x + 812, \text{ where } x = \delta\sqrt{\phi}, \tag{S2}$$

where $\delta$ is particle diameter (or the diameter of a sphere with equivalent volume) and $\phi$ is void volume fraction of the packing. Our unbiased estimator using softer deformable spheres gives an NRMSE of 0.0405 for particle surface area and 8.24 for open surface area. The latter result suggests that Equation S2 is relatively less generalizable to softer materials possibly because particle packing of softer materials results in less void space overall, contributing to lower open surface area and subsequent resolution issues given our crude approach for computing surface area. sp, spheres; ell, ellipsoids; rd, rods; ng, nuggets; def sp, deformable spheres; test a, unbiased estimator of softer deformable spheres with $\delta$ = 100, $\lambda$= 400, $\mu$ = 50. N = 10 domains (sp, def sp, test a) and N = 4 – 5 domains (ell, rd, ng).

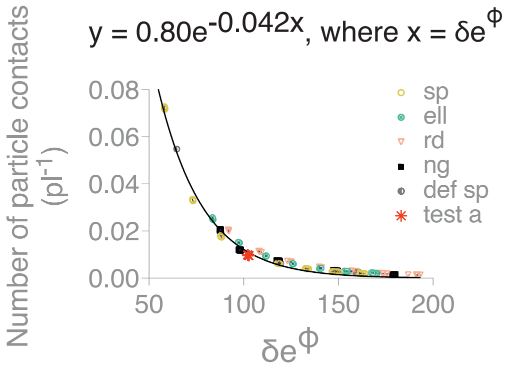


**Supplementary Figure 2 | Number of particle contacts.** Exponential decay equation ($R^2$ = 0.9869) for predicting the number of paths per picoliter as a function of the transformation:

$$y = 0.799e^{-0.0417x}, \text{ where } x = \delta e^{\phi}, \tag{S3}$$

where $\delta$ is particle diameter (or the diameter of a sphere with equivalent volume) and $\phi$ is void volume fraction of the packing. Our unbiased estimator using softer deformable spheres gives an NRMSE of 0.0203. sp, spheres; ell, ellipsoids; rd, rods; ng, nuggets; def sp, deformable spheres; test a, unbiased estimator of softer deformable spheres with $\delta$ = 100, $\lambda$= 400, $\mu$ = 50. N = 10 domains (sp, def sp, test a) and N = 4 – 5 domains (ell, rd, ng).

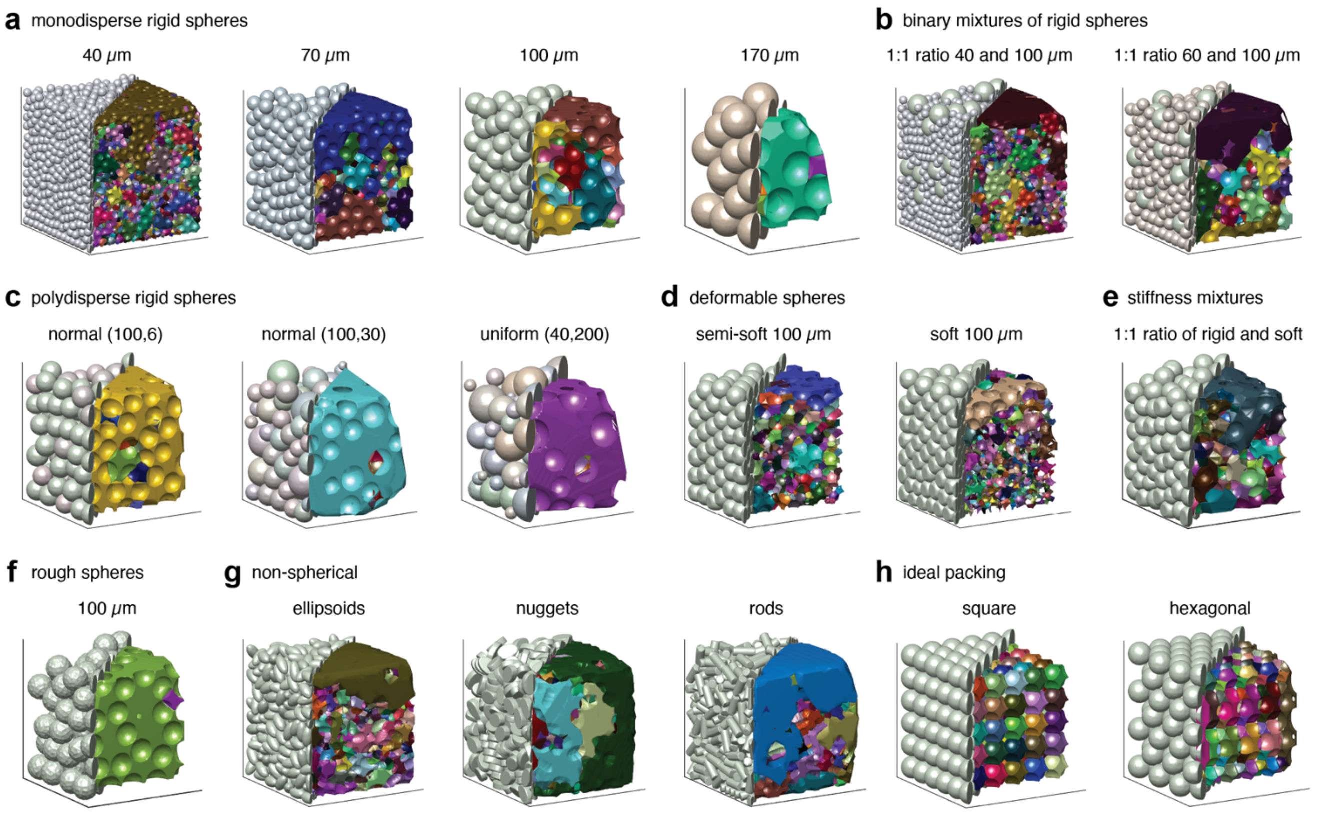


**Supplementary Figure 3 | Surface 3D pores for packings varying in particle size, shape, stiffness, and configuration.** Displaying simulated particle packings (left half) with corresponding surface 3D pores (right half). **a,** Labeling particle diameter. **b,** Labeling the ratio of particle diameters. **c,** (left) Normal distribution with mean, μ, of 100 μm and standard deviation, σ, of 6 μm; (middle) normal distribution with μ = 100 μm and σ = 30 μm; (right) uniform distribution with lower bound of 40 μm and upper bound of 200 μm. **d,** (left) Semi-soft spheres with Lamé parameters λ = 1,000, μ = 250; soft spheres with λ = 400, μ = 50. **e,** 100 μm spheres; soft particles described in d. **f,** Rough particles simulated by scattering 3 μm tall bumps along the surface of 100 μm spheres. **g,** Monodisperse and rigid with particle volume equal to the volume of a 100 μm diameter sphere. **h,** 100 μm diameter spheres.

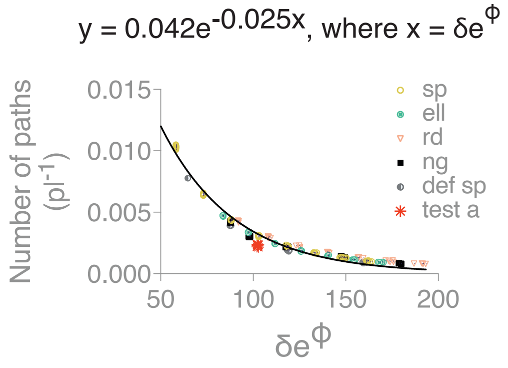


**Supplementary Figure 4 | Number of paths.** Empirically-derived exponential decay equation ($R^2$ = 0.9788) for predicting the number of paths per picoliter as a function of the transformation:

$$y = 0.0420e^{-0.0251x}, \text{ where } x = \delta e^{\phi}, \quad \text{(S4)}$$

where $\delta$ is particle diameter (or the diameter of a sphere with equivalent volume) and $\phi$ is void volume fraction of the packing. Our unbiased estimator using softer deformable spheres gives an NRMSE of 0.157. sp, spheres; ell, ellipsoids; rd, rods; ng, nuggets; def sp, deformable spheres; test a, unbiased estimator of softer deformable spheres with $\delta$ = 100, $\lambda$= 400, $\mu$ = 50. N = 10 domains (sp, def sp, test a) and N = 4 – 5 domains (ell, rd, ng).

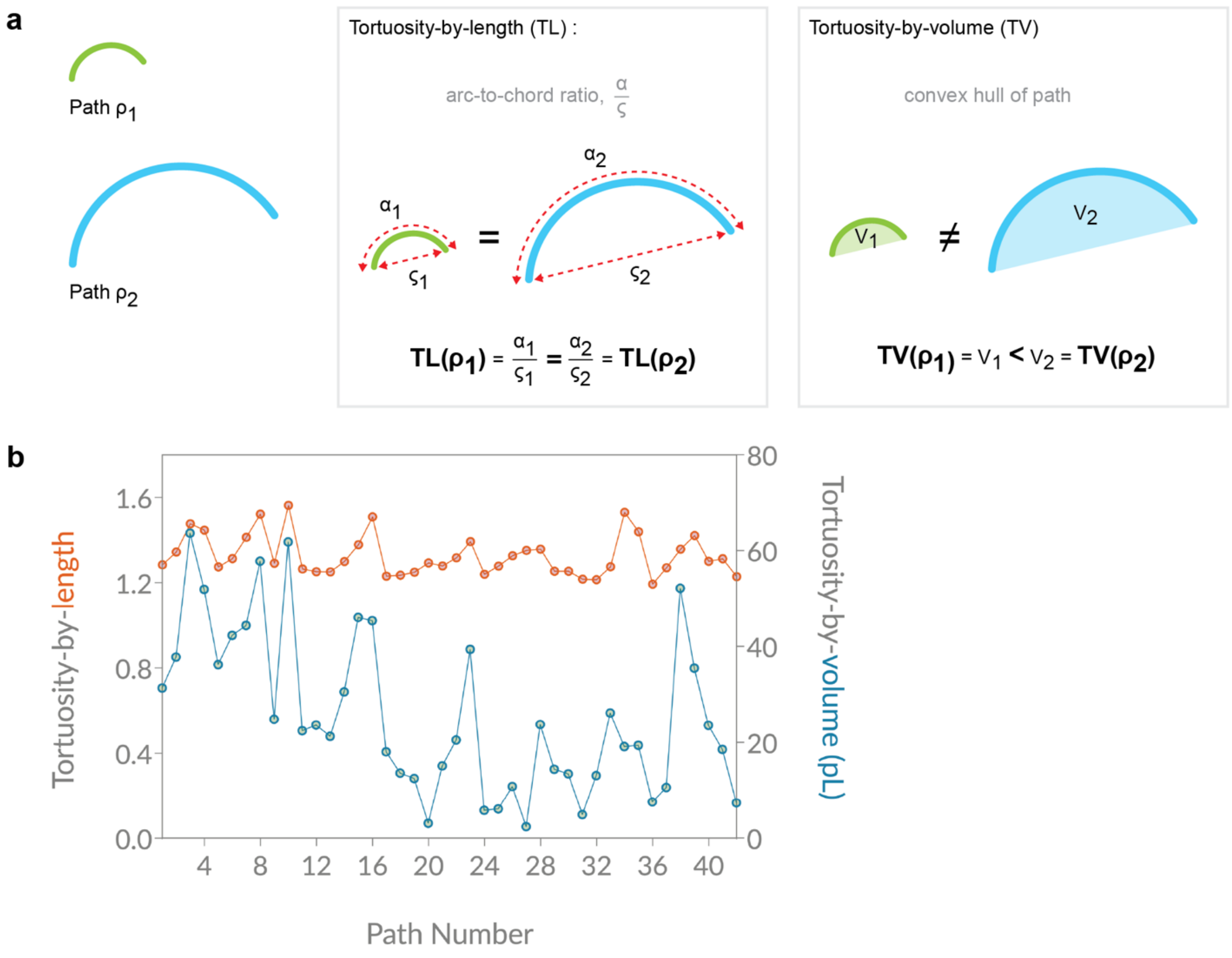


**Supplementary Figure 5 | Tortuosity-by-length (TL) and tortuosity-by-volume (TV). a,** TL is the path-to-chord ratio of a path, which is the length of the path divided by the distance between its endpoints. TV is the convex hull of a path, which is the smallest convex volume that encloses all points along the path. For two paths, $\rho_1 < \rho_2$, that share the same shape but differ in scale, TL($\rho_1$) = TL($\rho_2$), while TV($\rho_1$) < TL($\rho_2$). **b,** Data comparison of TL (orange) vs. TV (blue) for a simulated particle packing comprising monodisperse 130 µm diameter rigid spheres and containing 42 paths listed along the x-axis. A path is defined as the shortest distance along 1D ridges from the center to the surface (entrance door) of the scaffold. TV is measured in picoliters.

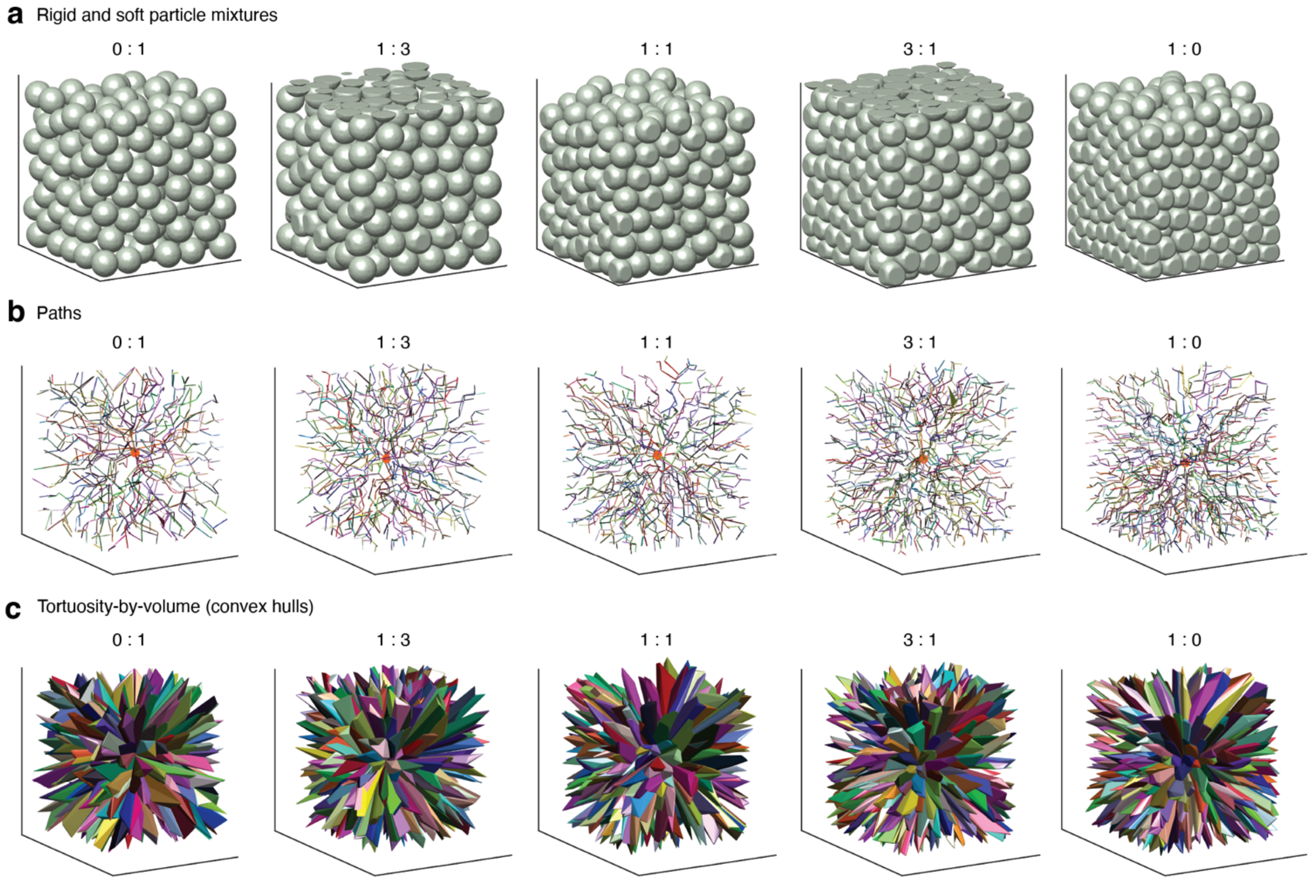


**Supplementary Figure 6 | Paths and tortuosity for randomly packed mixtures of rigid and deformable spheres. a,** Particle domains comprising mixtures of rigid and deformable (soft) 100μm diameter spheres. Ratios displayed above images. **b,** Displaying paths (along 1D-ridges), which radiate from the center of the packing toward entrance doors. **c,** Displaying convex hulls of paths, which are used to compute tortuosity-by-volume. Lamé parameters for soft particles: $\lambda = 400$, $\mu = 50$.

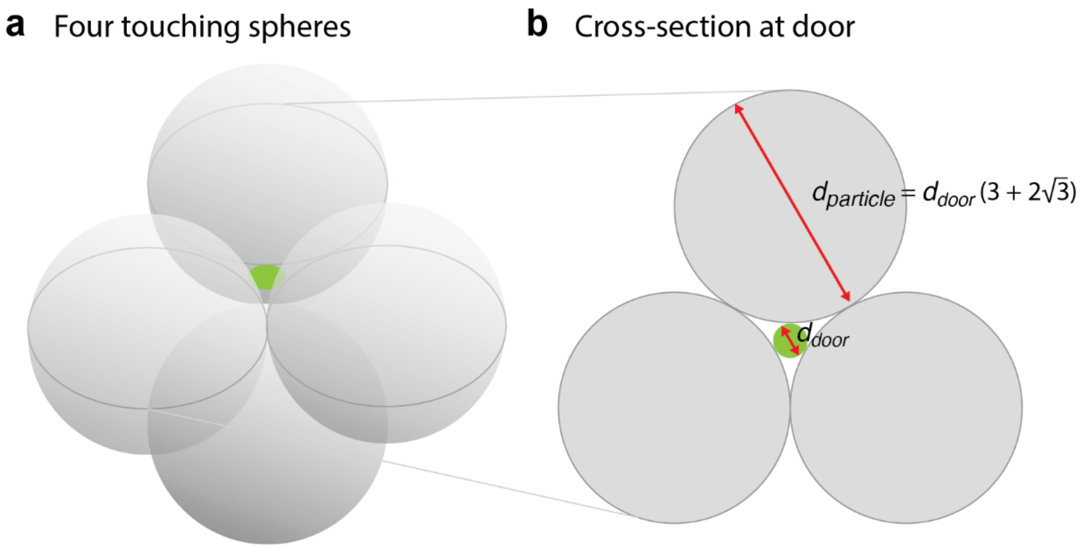


**Supplementary Figure 7 | Smallest pore type for packed rigid spheres. a,** Displaying four equal spheres configured in the tightest packing where each is in contact with the other. The void space within the touching particles forms the smallest pore type, bounded (in part) by four "doors" – one of which is shown in green. The green door lies within a plane defined by the center points of the three superior spheres. **b,** Displaying the cross-section along the plane of the door. The diameter of the door and the diameter of the circles are deterministically related according to the equation above.

Square packing

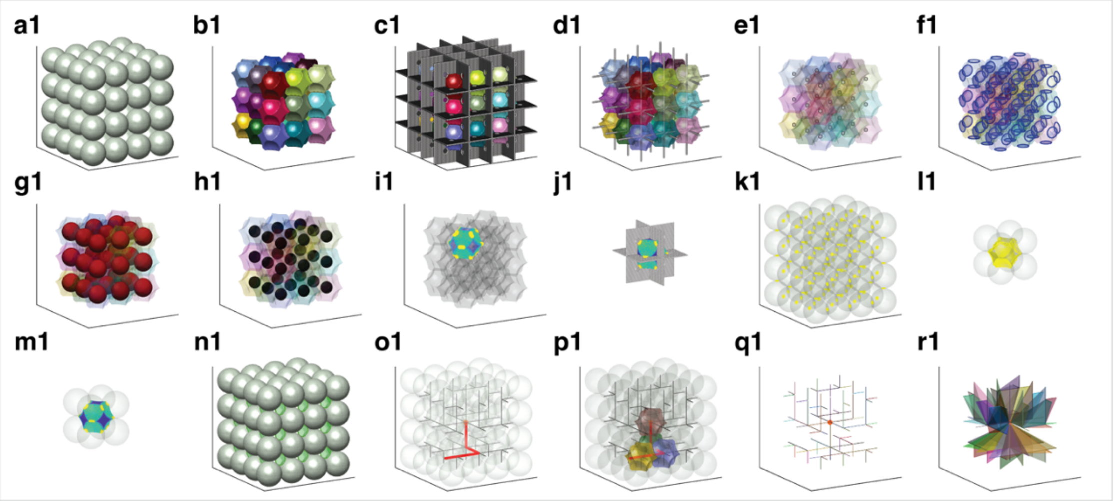


Hexagonal packing - Angle 1

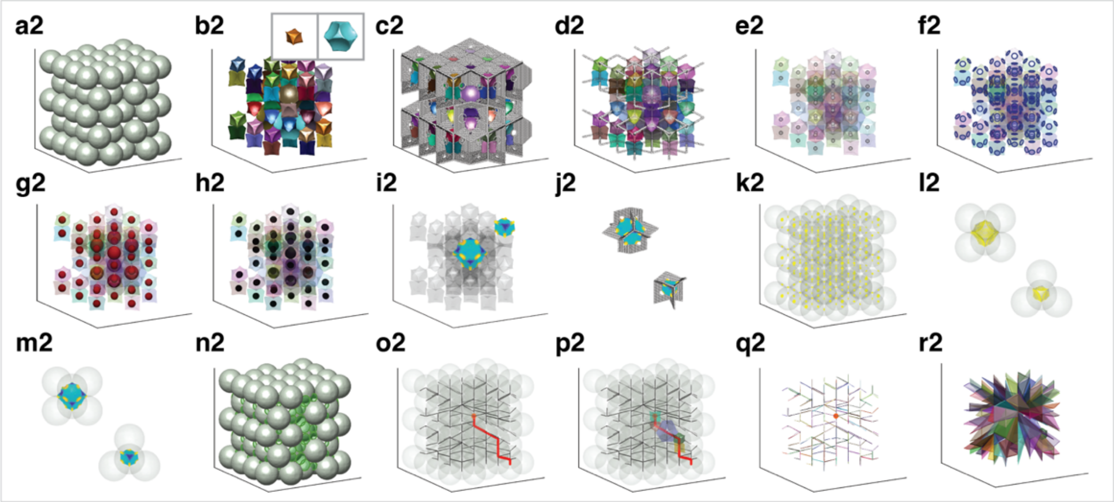


Hexagonal packing - Angle 2

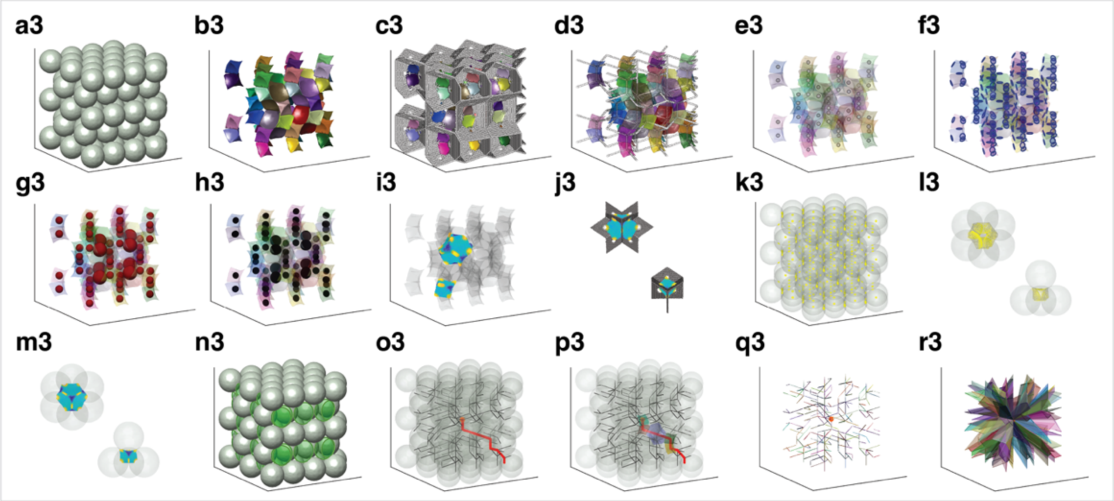

**Supplementary Figure 8 | Spatial landmarks of the medial axis and descriptors for square packing and hexagonal packing (shown from two different angles). a1-3,** Square and hexagonal packings of rigid spheres. **b1-3,** Interior 3D pores, with hexagonal packing containing two repeating pore types (b2 inset). **c1-3,** Displaying 2D ridges amongst interior pores. **d1-3,** Displaying 1D ridges (hallways) amongst interior pores. **e1-3,** Displaying peaks amongst interior pores. **f1-3,** Displaying interior doors (blue circles) amongst interior pores. **g1-3,** Displaying the largest enclosed sphere of each interior pore. **h1-3,** Displaying the PCA ellipsoid of each interior pore. **i1-3,** Highlighting pore vertices in yellow for sample pores. **j1-3,** Pore vertices (yellow) correspond to crawl spaces that lie along 2D ridges, shown for sample pores. **k1-3,** Displaying ligand hotspots (yellow) amongst particles. **l1-3,** Displaying accessible ligands (yellow) within sample pores that are surrounded by their enclosing particles. **m1-3,** Displaying the ligand concentration heatmap for sample pores that are surrounded by their enclosing particles. **n1-3,** Displaying entrance (or exit) doors (green circles) amongst particles. **o1-3,** Displaying paths (dark grey) amongst particles, highlighting a sample path in red. **p1-3,** Displaying the pores traversed by a sample path in red, with overlaying particles. **q1-3,** Displaying paths in isolation. **r1-3,** Displaying the convex hull of paths, used to compute tortuosity-by-volume (pL).

### Supplementary Notes

1. We derive the mean local thickness of a square, $MLT_s$, with length $\delta$:

   $$MLT_S(\delta) = \frac{8+\pi}{12} * \delta \approx (0.93 * \delta) < \delta.$$

   Indeed, we find that the $MLT_s$ of a square is less than its length.

2. For a pore, “vertices” and “crawl spaces” represent distinct concepts. In practice, however, both refer to the particle-particle contacts that produce the 2D ridges exiting the pore. Consequently, the total number of vertices and total number of crawl spaces are equivalent measurements.
